# A cellular and regulatory map of the GABAergic nervous system of *C. elegans*

**DOI:** 10.1101/052431

**Authors:** Marie Gendrel, Emily G. Atlas, Oliver Hobert

## Abstract

Neurotransmitter maps are important complements to anatomical maps and represent an invaluable resource to understand how signals are transmitted throughout the nervous system and how a nervous system is developmentally patterned. We report here a comprehensive map of neurons in the *C. elegans* nervous system that contain the neurotransmitter GABA, revealing twice as many GABA-positive neuron classes as previously reported. We define previously unknown glia-like cells that reuptake GABA, as well as “GABA reuptake neurons” which do not synthesize GABA but take it up from the extracellular environment. We used the map of GABA-positive neurons for a comprehensive analysis of transcriptional regulators that define the GABA phenotype. We synthesize our findings of specification of GABAergic neurons with previous reports on the specification of glutamatergic and cholinergic neurons into a nervous system-wide regulatory map which defines neurotransmitter specification mechanisms for more than half of all neuron classes in *C. elegans*.

## INTRODUCTION

Since the days of Ramon y Cajal, the generation of maps of the brain constitutes a central pursuit in the neurosciences. The nervous system of the nematode *C. elegans* constitutes the currently best mapped nervous system. Available *C. elegans* nervous system maps include a lineage map of all neurons (Sulston, 1983) and an anatomical map that describes all individual neuron types not just in terms of overall morphology but also synaptic connectivity (Jarrell et al., 2012; White et al., 1986). One type of map that complements anatomical maps and that is critical to understand neuronal communication is a map that assigns neurotransmitter identity to all neurons in the nervous system. Comprehensive maps of modulatory, monoaminergic neurons (i.e. serotonergic, dopaminergic) have been known for some time in *C. elegans* (Chase and Koelle, 2007), but comprehensive maps of the most prominent fast-acting acting neurotransmitter systems employed throughout all animal nervous systems – glutamate (Glu), acetylcholine (ACh) and GABA - are only now emerging. We have recently defined the complete set of glutamatergic (Serrano-Saiz et al., 2013) and cholinergic neurons in *C. elegans* (Pereira et al., 2015) and in this third and last paper of a trilogy of neurotransmitter-mapping papers, we describe our analysis of GABA-positive neurons, expanding previous work that had begun to define GABAergic neurons in *C. elegans* (McIntire et al., 1993b).

GABA is a fast acting neurotransmitter that is broadly used throughout all vertebrate and invertebrate nervous systems. In vertebrates, GABA is used as neurotransmitter by many distinct neuron types throughout the CNS (30-40% of all synapses are thought to be GABAergic; (Docherty et al., 1985)) and alterations of GABAergic neurotransmission are the cause of a number of neurological diseases in humans (Webster, 2001). One intriguing issue, unresolved in vertebrates due to the complexity of their nervous systems, is the cellular source of GABA and the fate of GABA after cellular release. The expression of the biosynthetic enzyme for GABA, glutamic acid decarboxylase (GAD), defines neurons that have the capacity to synthesize GABA, but the existence of plasma membrane transporters for GABA (called GAT) indicates that GABA can also be “acquired” by neurons via transport and not synthesis (Zhou and Danbolt, 2013). Does GABA reuptake merely occur to clear GABA, thereby controlling the duration of a GABAergic signal, or do cells take up GABA to then reemploy it, e.g. by using vesicular GABA transporters (VGAT) to synaptically release GABA? A precise map of GAD-, GAT-and VGAT-expressing neurons with single neuron resolution would shed light on these issues, but has not yet been produced in vertebrate nervous systems. In this resource paper we provide such a map in the nematode *C. elegans*.

Previous studies have ascribed a GABAergic neurotransmitter identity to 26 *C. elegans* neurons, which fall into six anatomically and functionally diverse neuron classes. These numbers amount to less than 10% of all neurons (302 hermaphroditic neurons) and neuron classes (118 anatomically defined neuron classes)(McIntire et al., 1993a; McIntire et al., 1993b; Schuske et al., 2004). Not only is this substantially less than the number of neurons that use conventional excitatory neurotransmitters (Glu: 39 classes, ACh: 52 classes; (Pereira et al., 2015; Serrano-Saiz et al., 2013)), but, given the abundance of GABAergic interneurons in vertebrates, it is also striking that only one of the previously identified GABA neurons is an interneuron (McIntire et al., 1993b). However, the *C. elegans* genome contains at least seven predicted ionotropic GABA receptors (Hobert, 2013) and at least some of them are expressed in cells that are not synaptically connected to the previously defined GABA neurons (Beg and Jorgensen, 2003; Jobson et al., 2015). We therefore suspected that additional GABAergic neurons may have been left undetected. Using a refined GABA antibody staining protocol and improved reporter gene technology, we extended the original set of six GABA-positive neuron classes by another 10 additional GABA-positive cell types, eight of them neuronal cell types.

The knowledge of the complete and diverse set of neurons that share the expression of a specific neurotransmitter system allow one to ask how the expression of a shared identity feature is genetically programmed in distinct neuron types. As mentioned above, the usage of GABA as a neurotransmitter represents a unifying terminal identity feature for a diverse set of neurons in invertebrate and vertebrate nervous systems. Given the diversity of GABAergic neuron types, it is perhaps not surprising that no unifying theme of GABAergic identity specification has yet been discovered. Nevertheless, while distinct GABAergic neuron types use distinct transcription regulatory codes, some transcription factors appear to be repeatedly used by distinct GABAergic neuron types. For example, in vertebrates, the GATA-type transcription factors GATA2/3 is employed for GABAergic identity specification by midbrain and spinal cord neurons (Achim et al., 2014; Joshi et al., 2009; Kala et al., 2009; Lahti et al., 2016; Yang et al., 2010). We explore here whether the theme of reemployment of a transcription factor in different GABAergic neurons also exists in *C. elegans*.

Another question that pertains to the development of GABAergic neurons relates to the stage at which regulatory factors act to specify GABAergic neurons. Previous studies on the specification of vertebrate GABAergic neurons have so far uncovered factors that act at distinct stages of GABAergic neuron development (Achim et al., 2014). However, there is still a remarkable dearth of knowledge about transcriptional regulators that are expressed throughout the life of GABAergic neurons to not only initiate but also maintain the differentiated state of GABAergic neurons. Such type of late acting transcriptional regulators have previously been called “terminal selectors” and these terminal selectors have been identified to control the identity of distinct neuron types utilizing a variety of distinct neurotransmitter systems (Hobert, 2008, 2016a).

In *C. elegans*, previous work has shown that the Pitx2-type transcription factor *unc-30* selectively specifies the identity of D-type motor neurons along the ventral nerve cord (Cinar et al., 2005; Eastman et al., 1999; Jin et al., 1994). As a terminal selector of D-type motor neuron identity, UNC-30 protein directly controls the expression of GABA pathway genes (Eastman et al., 1999) as well as a plethora of other D-type motor neuron features (Cinar et al., 2005), including their synaptic connectivity (Howell et al., 2015). Since *unc-30* is expressed in non-GABAergic neurons (Jin et al., 1994), *unc-30* is not sufficient to induce GABAergic neuronal identity, possibly because *unc-30* may act with as yet unknown cofactors in the D-type motor neurons. We identify here a putative cofactor for *unc-30* in the form of the *elt-1* gene, the *C. elegans* ortholog of the vertebrate Gata2/3 transcription factors, which specify GABAergic neurons in vertebrates (Achim et al., 2014).

The acquisition of GABAergic identity of *C. elegans* neurons other than the D-type GABAergic motor neurons was less well understood. The RIS, AVL and DVB neurons display differentiation defects in animals lacking the *lim-6* homeobox gene, the *C. elegans* ortholog of the vertebrate *Lmx1* LIM homeobox gene (Hobert et al., 1999; Tsalik et al., 2003) and the RME neurons display differentiation defects in animals that carry a mutation in the *nhr-67* orphan nuclear hormone receptor, the *C. elegans* ortholog of vertebrate Tlx/NR2E1 gene and the fly gene Tailless (Sarin et al., 2009) or in animals lacking *ceh-10*, the *C. elegans* ortholog of the vertebrate *Vsx/Chx10* Prd-type homeobox gene (Forrester et al., 1998; Huang et al., 2004). However, the extent of the differentiation defects in these distinct mutant backgrounds have not been examined in detail. We show here that *nhr-67* controls all GABAergic identity features in the AVL, RIS and RMEL/R and RMED/V motorneurons, where it collaborates with distinct homeobox genes, *lim-6* in AVL and RIS, *ceh-10* in RMED and *tab-1*, the *C. elegans* ortholog of vertebrate Bsx, in RMEL/R. We identify additional homeobox genes that control the identity of GABAergic neurons that we newly identify here. Taken together, our systems-wide analysis of GABA-positive cells in *C. elegans* identifies a number of distinct GABA-positive cell types that acquire and utilize GABA via diverse mechanisms and provides an extensive picture for the specification of this critical class of neurons.

## RESULTS

### Identifying GABAergic neurons in the *C. elegans* nervous system

The previously reported set of GABAergic neurons (six neuron classes: RME, AVL, RIS, DVB, DD, VD; **Table 1**) were defined by anti-GABA antibody staining and expression of reporter transgenes that monitor expression of genes that encode the GABA biosynthetic enzyme glutamic acid decarboxylase (GAD/UNC-25) and the vesicular GABA transporter (VGAT/UNC-47)(Jin et al., 1999; McIntire et al., 1993b; McIntire et al., 1997). Using a modified GABA staining protocol, we observed the presence of GABA in the six previously described GABAergic neuron classes RME, AVL, RIS, VD, DD and DVB, but also detected staining in additional set of eight neuronal cell types that stain for GABA (RIB, SMDD, SMDV, AVB, AVA, AVJ, ALA, AVF; **Table 1**, **Fig.1**). The identity of these GABA-positive neurons was confirmed by GABA-staining of transgenic animals that express cell-type specific markers (data not shown). Staining of these newly identified GABA-positive cells is generally weaker than in the previously identified GABA neurons (**Fig.1**). In the vertebrate CNS, distinct GABAergic neuron types also show different levels of anti-GABA staining (J. Huang, pers.comm.). Anti-GABA staining in these newly identified cells is abolished in animal lacking the gene that codes for the enzymes that synthesizes GABA, *unc-25/GAD*, corroborating that the staining indeed reports on the presence of GABA (**Fig.1**). The same pattern of staining is observed in early larvae and adult worm, with the sole exception of the AVJ neuron pair which stains more strongly in larval compared to adult stages.

**Fig.1:**
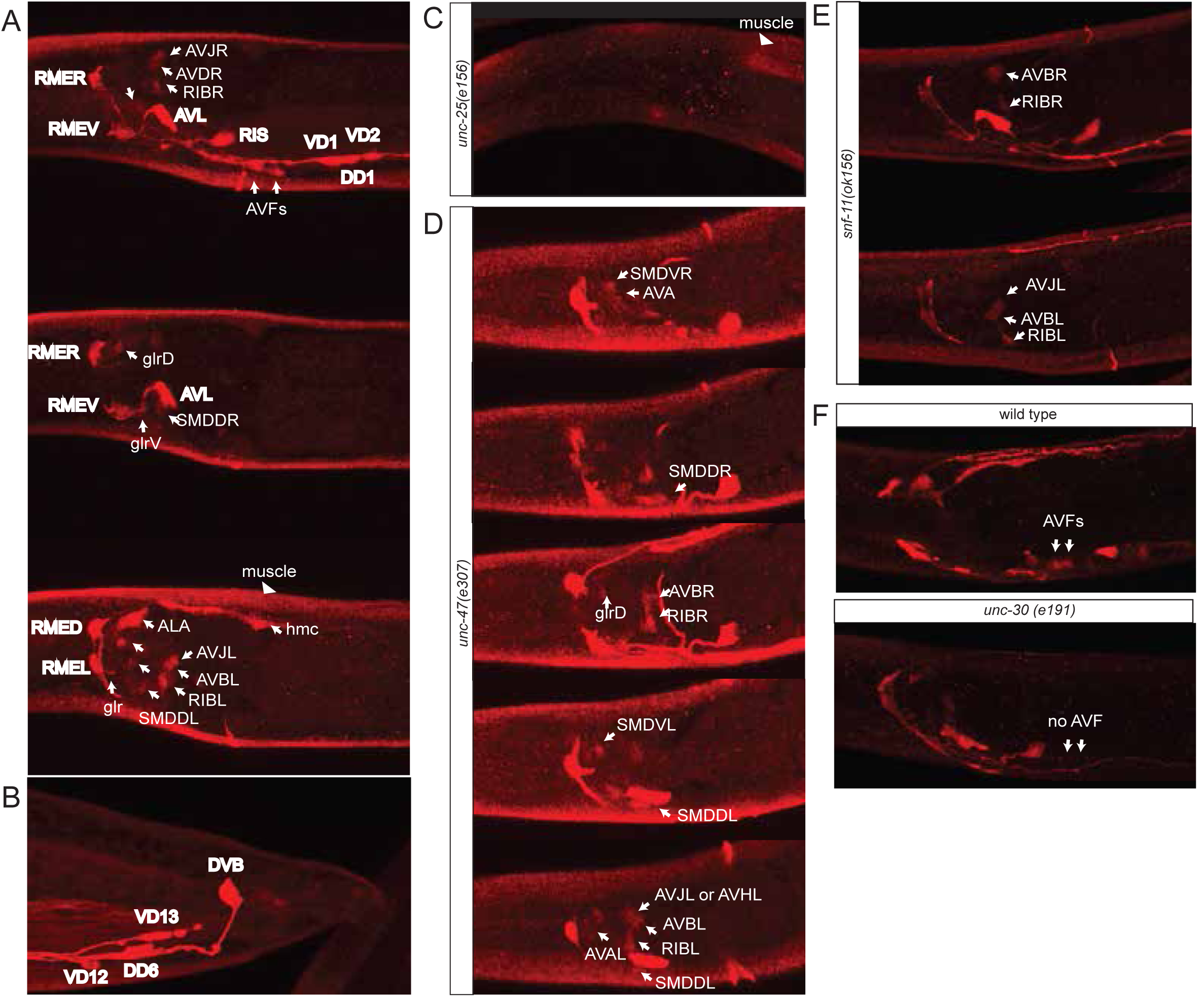
Anti-GABA staining defines the GABAergic nervous system of *C. elegans*. **A:** Three different optical sections of the head of a single adult hermaphrodite. Previously identified GABA neurons (McIntire et al., 1993b) are shown in bold. **B:** Tail of an adult hermaphrodite. **C:** Head of an adult *unc-25/GAD(-)* hermaphrodite. Absence of GABA staining illustrating that GABA staining is specific. Staining in muscles is diminished but not eliminated completely. **D:** Five different optical sections of the head of a single adult *unc-47/VGAT(-)* hermaphrodite, which illustrates the dependence of some GABA-positive cells on vesicular GABA secretion. Only the newly identified GABA neurons are labeled. SMD is more strongly stained in *unc-47* mutants compared to wild type animals where it is weaker and more variable. **E:** Five different optical sections of the head of of a single adult *snf-11/GAT(-)* hermaphrodite, which illustrates the dependence of some GABA-positive cells on GABA reuptake. Only the newly identified GABA neurons are labeled. **F:** AVF neuron staining is absent in *unc-30* mutant animals.

**Table 1:**
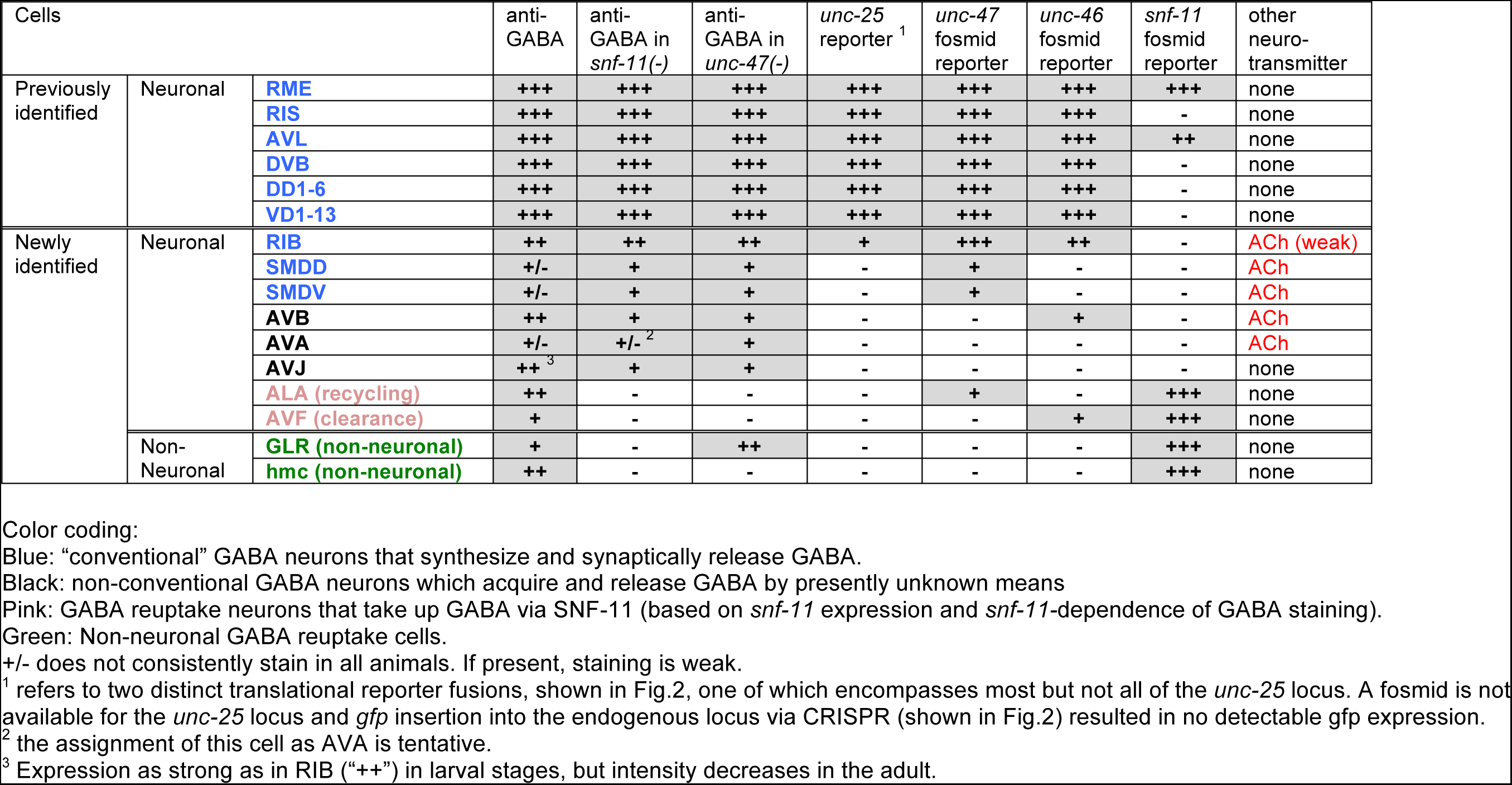
GABA-positive cells

### GABA reuptake neurons

Neurons that package a specific neurotransmitter and release it to signal to downstream neurons do not necessarily synthesize this neurotransmitter, but may rather internalize it from their environment via neurotransmitter-specific reuptake systems. For example, spinal cord motor neurons in the rat do not synthesize GABA but reuptake GABA after release from presynaptic neurons (Snow et al., 1992). GABA synthesis is usually examined by analyzing the expression pattern of the GABA-synthesizing enzyme glutamic acid decarboxylase (GAD), encoded by the *unc-25* gene in *C. elegans*. A previously published reporter transgene for the *unc-25* locus shows expression in the set of previously identified GABA neurons (Jin et al., 1999). We also detect expression of this reporter in the newly identified GABA-positive RIB interneuron pair but not in any of the other newly identified GABA-positive neurons (**Fig.2**). This may be due to the previous reporter only containing parts of the *unc-25* locus (e.g. missing the last few intron; **Fig.2B**). No fosmid was available to generate a fosmid-based reporter gene construct, and *mNeonGreen* insertion into the *unc-25* locus using the CRISPR/Cas9 system (**Fig.2B**) did not produce any detectable expression. Since the expression pattern analysis of *unc-25/*GAD therefore remains inconclusive, we chose alternative ways to assess whether the remaining, newly identified GABA-positive neurons synthesize their own GABA or reuptake of GABA from other cells. Specifically, we performed three sets of experiments: (a) we GABA-stained animals that are unable to release GABA because they lack the GABA vesicular transporter VGAT/SLC32, called UNC-47 in *C. elegans* (McIntire et al., 1997); (b) we anti-GABA-stained animals that lack the sole ortholog of the GABA reuptake transporter GAT/SLC6A1, called SNF-11 in *C. elegans* (Jiang et al., 2005; Mullen et al., 2006); and (c) we analyzed the expression pattern of SNF-11/GAT. We find that GABA staining of the ALA and AVF neurons is abolished in either *unc-47* or *snf-11* mutants, suggesting that these neurons do not synthesize their own GABA, but obtain synaptically released GABA by reuptake via the plasma membrane transporter SNF-11/GAT (**Fig.1**). In support of this notion, both ALA and AVF express a fosmid-based reporter that we generated for the *snf-11* locus (**Fig.2**). We therefore termed the AVF and ALA neurons “GABA reuptake neurons”. Within the nervous system, the *snf-11* fosmid-based reporter is also expressed in the previously characterized, GABA-synthesizing RME and AVL neurons, but not the other “classic” GABA-synthesizing RIS, DVB or D-type neurons (**Fig.2**).

**Fig.2:**
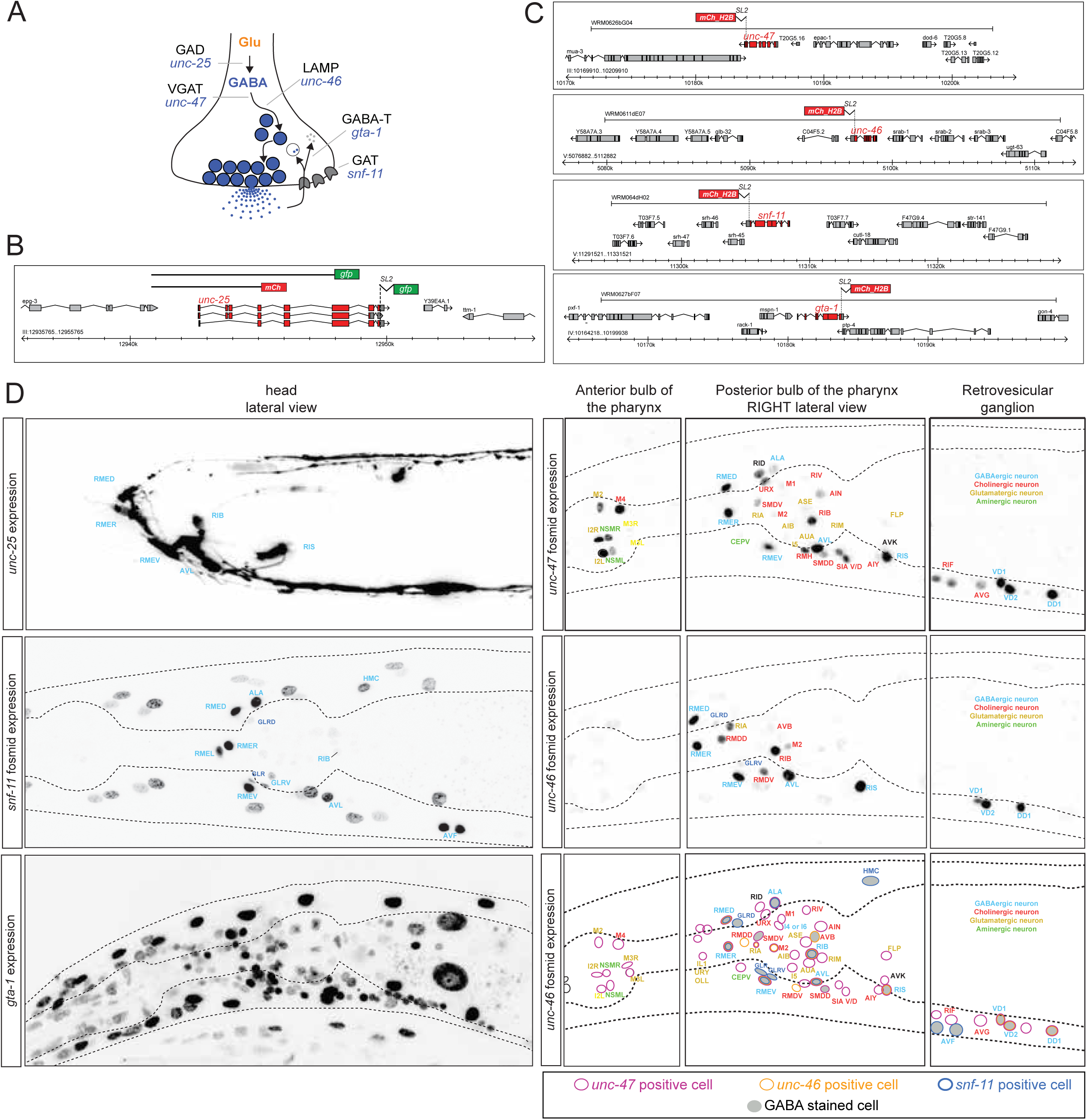
Reporter gene constructs recapitulate GABA antibody staining in the nervous system of the *C. elegans* hermaphrodite. **A:** GABA synthesis, secretion, reuptake and degradation pathway. **B:** *unc-25* transcriptional reporter transgene schematics. **C:** Fosmid-based reporter transgenes for *unc-46, unc-47, snf-11* and *gta-1*. Note that reporters are targeted to the nucleus. **D:** Fosmid reporter expression. Images are color-inverted. The last panel shows a summary of all reporter expressing neurons.

Where could AVF and ALA receive GABA from? According to the *C. elegans* connectivity map (White et al., 1986), the unpaired ALA neuron is not a direct postsynaptic target of any GABA-positive neurons. However, as assessed by the examination of electron micrographs produced by White et al., the processes of ALA are directly adjacent to the newly identified GABAergic SMD neurons (White et al., 1986)(S. Cook, pers.comm.). This indicates that ALA may absorb GABA released by the SMD neurons, thereby modulating GABA transmission between SMD and its target neurons. Consistent with this notion, we find that prevention of GABA release in *unc-47/VGAT* mutants results in a considerable increase in GABA staining in the SMDD and SMDV neurons (**Fig.1**).

The AVF reuptake neuron class is located at a different position in the *C. elegans* head ganglia and display projections very distinct from the ALA neuron. It extends a process into the ventral nerve cord (http://wormatlas.org/neurons/Individual%20Neurons/AVFframeset.html) and may take up GABA from proximal GABAergic D-type motor neurons which populate the nerve cord. To test this notion experimentally, we removed GABA selectively from D-type motor neuron, using the *unc-30* mutant strain in which D-type motor neurons fail to produce GABA (Jin et al., 1994). In these animals GABA staining in AVF is abolished (**Fig.1F**), suggesting that AVF indeed absorbs GABA from the D-type motor neurons. The AVF neurons also receive a single synapse from the GABAergic AVL neuron (White et al., 1986), suggesting another possible source of GABA.

### VGAT expression suggests the existence of “recycling neurons” and “clearance neurons”

We next sought to assess which of the GABA-positive neurons have the ability to synaptically release GABA via the canonical vesicular GABA transporter VGAT/UNC-47 (McIntire et al., 1997). To this end, we generated a fosmid-based *unc-47/VGAT* reporter construct which contains considerably more sequence information than previous reporter constructs. This reporter shows a much broader pattern of expression compared to the original reporter (McIntire et al., 1997); it is expressed in the original set of GABAergic neurons and also in the newly identified GABA-positive ALA, RIB, SMDD and SMDV neurons, indicating that these neurons not only contain GABA but can also synaptically release it (**Table 1**; **Fig.2**).

We could not detect *unc-47* fosmid reporter expression in the AVF neurons, which do not synthesize but only reuptake GABA, indicating that these neuron may only function to clear, but not re-release GABA. We cannot exclude the possibility that GABA may be released by non-conventional mechanisms. We also could not detect *unc-47/VGAT* in the GABA-positive AVA, AVB and AVJ neurons. Even though the absence of proper reagents to assess UNC-25/GAD expression prevents us from proving this point, we surmise that these neurons may either synthesize their own GABA (since their positive GABA-positive nature does not depend on the SNF-11 reuptake transporter (**Table 1**, **Fig.1**)) or they may acquire GABA via gap junctions that these neurons make with other GABA-positive neurons (neurotransmitter diffusion via gap junctions has been observed in vertebrates and called “neurotransmitter coupling”; (Vaney et al., 1998)). However, the absence of *unc-47/VGAT* expression in these neurons suggests that these neurons may use non-conventional and presently not understood GABA release mechanisms, which are also thought to exist in the vertebrate CNS (Koch and Magnusson, 2009).

Beyond the above-mentioned GABA(+) cells, the *unc-47* reporter fosmid is expressed in a substantial number of additional neurons (**Fig.2**, summarized in **Table 2**). Since those neurons show no evidence for GABA expression, we suspect that in these neurons *unc-47/VGAT* may transport an as yet unknown neurotransmitter.

We also generated a fosmid-based reporter for the *unc-46* locus, which encodes a LAMP-like protein, previously reported to be important for UNC-47/VGAT localization (Schuske et al., 2007). This fosmid-based reporter is expressed in most neurons that are GABA(+) and UNC-47(+)(**Table 1**; **Fig.1**, **Table 2**). Interestingly, neurons that do not contain GABA but express *unc-47* mostly do not express *unc-46* (**Table 2**). This suggests that the non-GABA-related functions of *unc-47* may not strictly require *unc-46*.

**Table 2:** Summary of *unc-47/VGAT* and *unc-46/LAMP-*expressing neurons in GABA-negative neurons. Expression in GABA-positive neurons is shown in Table 1. Pharyngeal neurons are separated by an double line. +/-, + and ++ represent relative expression levels.

| Cells | anti-GABA | *unc-47*<br>fosmid<br>reporter | *unc-46*<br>fosmid<br>reporter | other neuro-<br>transmitter |
| --- | --- | --- | --- | --- |
| AIB | - | +/- | - | Glu |
| AIN | - | ++ | - | Glu |
| AIY | - | + | - | ACh |
| ALN | - | + | - | ACh |
| ASE | - | + | - | Glu |
| AUA | - | +/- | - | none |
| AVG | - | + | - | none |
| AVK | - | + | - | Glu |
| CEPV | - | + | - | dopamine |
| DVA | - | ++ | - | ACh |
| FLP | - | + | - | Glu |
| IL1 | - | + | - | Glu |
| OLL | - | +/- | - | Glu |
| PLN | - | + | - | ACh |
| RIA | - | +/- | ++ | Glu |
| RID | - | ++ | - | none |
| RIF | - | ++ | - | ACh |
| RIM | - | +/- | - | Glu |
| RIV | - | + | - | ACh |
| RMDD/V | - | - | ++ | ACh |
| SIA | - | ++ | - | ACh |
| URX | - | ++ | - | ACh |
| URY | - | +/- | - | Glu |
| MI | - | ++ | - | Glu |
| M1 | - | + | - | ACh |
| M2 | - | + | + | ACh |
| M3 | - | + | - | ACh |
| M4 | - | ++ | - | ACh |
| M5 | - | + | - | ACh/Glu |
| I2 | - | ++ | - | Glu |
| I4 or I6 | - | ++ | - | none |
| I5 | - | + | - | Glu |
| NSM | - | + | - | 5HT |

Lastly, we examined whether the expression of the enzyme that degrades GABA, GABA transaminase (GABAT), is expressed or perhaps even enriched in the GABA clearance neurons. *C. elegans* contains a single ortholog of GABAT, which we termed *gta-1*. We find a fosmid-based reporter of *gta-1* to be ubiquitously expressed (**Fig.2**), mirroring the very broad tissue distribution of vertebrate GABAT and consistent with the notion that GABAT uses substrates other than GABA (Jeremiah and Povey, 1981).

In conclusion, we have added another eight GABA-positive neuron types to the previous list of six GABA-positive neuron types (**Table 1**). Two of these neuron types, the RIB neurons and the SMDD/V neuron types are “conventional” GABA neurons similar to the previously characterized GABA-positive neurons, in that they likely synthesize and synaptically release GABA. Two neuron types (ALA and AVF) are GABA reuptake neurons that acquire GABA from neighboring cells to either simply remove GABA (AVF) or possibly also reuse GABA (ALA). Three other neuron types (AVA, AVB and AVJ), two of them previously shown to be cholinergic (AVA, AVB; (Pereira et al., 2015)) acquire GABA by as yet unknown means; whether and how they employ GABA for signaling is also presently not clear.

### GABA in non-neuronal cells

In addition to neuronal staining, we also detected GABA in three classes of non-neuronal cells, the head mesodermal cell (hmc), the glia-like GLR cells and body wall muscle (**Fig.1**). Expression in all these cells depends on *unc-25/*GAD (**Fig.1**; note that residual staining in muscle remains in body wall muscle). Body wall muscle may take up GABA after release by the D-type motorneurons which innervate body wall muscle. Consistent with this notion, *snf-11* is expressed in body wall muscle (**Fig.2**)(Mullen et al., 2006).

The hmc is an intriguing cell with no previously ascribed function. It is located above the posterior bulb of the pharynx and extends processes along the ventral and dorsal nerve cord (http://www.wormatlas.org/ver1/handbook/mesodermal.htm/hmc.htm). The processes of the hmc are in proximity to the processes of a number of GABAergic neurons (RMED/V and D-type motor neurons)(Hall and Altun, 2007) and therefore in the proper place to clear GABA. We indeed find that hmc expresses the GABA reuptake transporter *snf-11/GAT* (**Fig.2D**) and that GABA staining of the hmc is abolished in *snf-11/GAT* mutants (**Fig.1E**). We also find that GABA staining in the hmc is reduced in *unc-30* mutants (in which D-type motor neurons fail to be specified; (Jin et al., 1994)), indicating that the source of GABA in the hmc are indeed the D-type motor neurons. Since we do not detect *unc-47/GAT* expression in the hmc, the hmc likely operates as a GABA clearance cell, like the AVF neurons.

The glia-like GLR cells are another intriguing GABA-positive cell type. GLR cells, which have no assigned function yet, are located directly adjacent to the nerve ring (http://www.wormatlas.org/ver1/handbook/mesodermal.htm/glr.htm). Each GLR cell extends a thin, sheet-like process that lies inside the nerve ring. Like the AVF and ALA neurons and the non-neuronal hmc, the GLR cells express *snf-11/GAT* (**Fig.2**) and GABA-staining is strongly reduced in *snf-11/GAT* mutants (**Fig.1**). Curiously, staining is still observed in *unc-47/VGAT* mutants. In these mutants, GABA may accumulate in RME neurons which are heavily gap-junctioned the GLR glia cells (White et al., 1986) and GABA may pass through these gap junctions. The passive transfer of neurotransmitters through gap junctions has been termed “neurotransmitter coupling” and occurs, for example, in the amacrine and bipolar cells of the vertebrate retina (Vaney et al., 1998).

### GABAergic neurons in the male nervous system

In males, we observed the same set of GABA-positive neurons as observed in hermaphrodites. In addition, GABA staining is observed in one prominent class of male-specific interneurons, the EF neurons, which had previously no neurotransmitter assigned (**Fig.3**). The EF1 and EF2 interneurons are located in the dorsorectal ganglion, while EF3 (and the rarely generated EF4) is in the preanal ganglion (**Fig.3**). The EF neurons are so-called “type II interneurons”, which relay sensory information from male-specific tail sensory structures into the sex-shared nervous system in the head (Jarrell et al., 2012). All the EF neurons also express *unc-25/GAD, unc-47/VGAT* and *snf-11/*GAT, demonstrating that these neurons synthesize, release and reuptake GABA (**Fig.3**). Apart from the EF neurons, three additional male-specific sensory neurons stain with anti-GABA antibodies, the R2A, R6A and R9B pairs of ray neurons (**Fig.3**).

**Fig.3.**
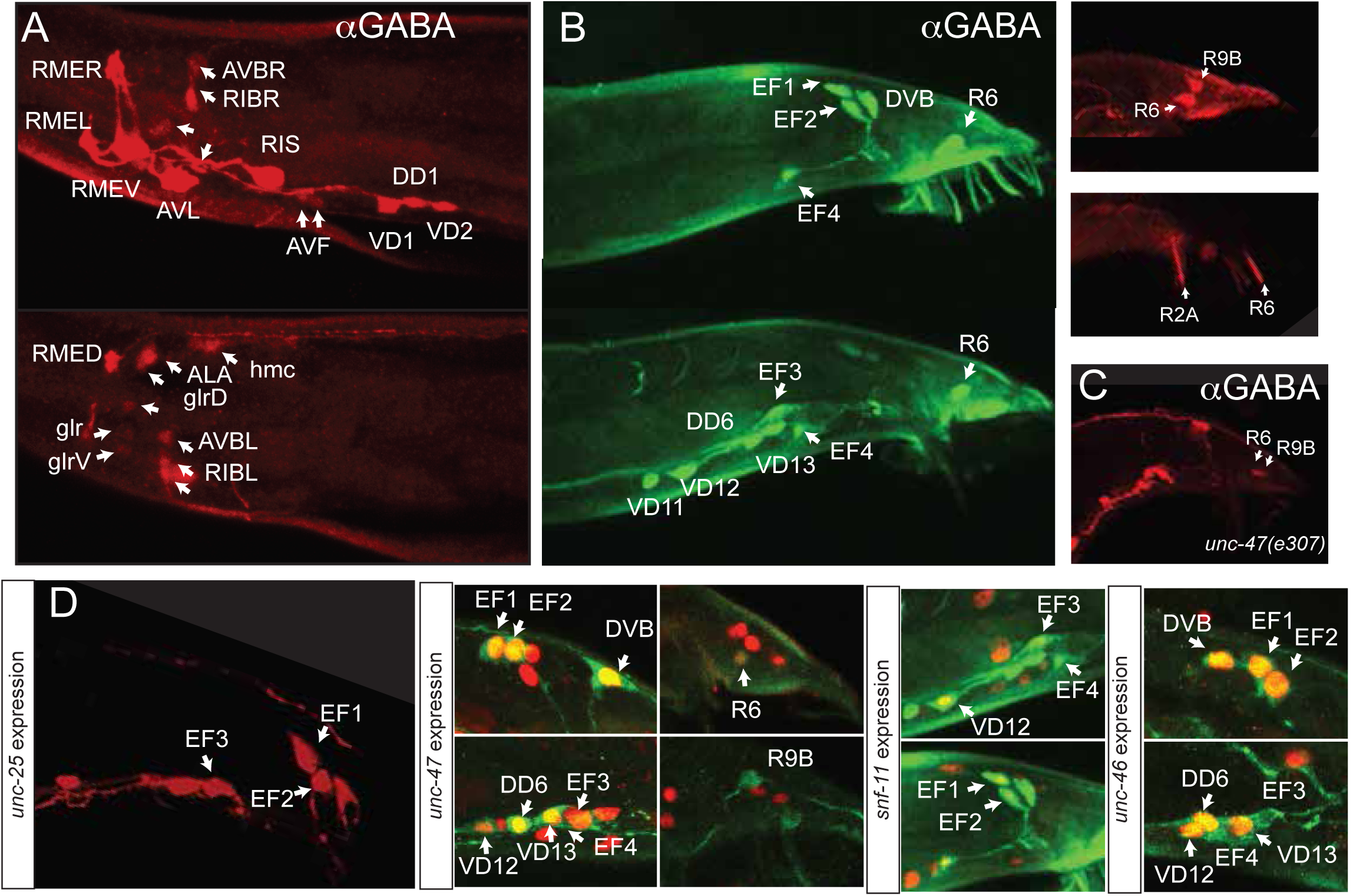
GABAergic neurons in the male. **A-C:** Anti-GABA staining of adult males in the head (A; two optical sections; B: two optical section with green fluorescent secondary antibody in left two panels (ray neuron staining is background) and red fluorescent secondary antibody in upper right panels (ray neuron staining has no background; C: in *unc-47* mutant animals). **D:** Reporter gene expression in adult male tail. Left panel: *unc-25* reporter, other panel show red fosmid reporter (nucleus) and green GABA staining.

Lastly, we noted an intriguing sexual dimorphism of *snf-11* expression in the VD12 neuron, which is generated in both sexes and produces GABA in both sexes, based on *unc-25/GAD* expression and GABA staining. In hermaphrodites, none of the 13 VD neurons express *snf-11/GAT*, but in males, VD12 expresses *snf-11*(**Fig.3**). While VD12 receives a multitude of male-specific synaptic inputs (Jarrell et al., 2012), it is not directly postsynaptic to GABA-positive male tail neurons (i.e. EF or R6B). Perhaps it is particularly relevant to limit the amount of GABA released by VD12 onto muscle in the male, but not hermaphrodite tail.

### Synaptic connectivity of GABA-positive neurons

We used the synaptic wiring diagram of the hermaphrodite elucidated by John White and colleagues (Varshney et al., 2011; White et al., 1986)(www.wormwiring.org) to assess the extent of GABAergic synaptic innervation throughout the entire nervous system. Of the 118 neuron classes of the hermaphrodite, 47 neuron classes are innervated by GABA(+); UNC-47(+) neurons (**Table 3**). The RIB and SMD neurons are the GABAergic neurons with the most synaptic partners (**Table 3**). Both neurons also employ ACh as a neurotransmitter, even though in each case, one neurotransmitter system appears to predominate: The RIB neurons express barely detectable levels of VAChT/ChAT (Pereira et al., 2015), while their GABA staining is easily detectable. Vice versa, the SMD neurons strongly express VAChT/ChAT (Pereira et al., 2015), but their GABA staining is weaker and more variable.

**Table 3:**
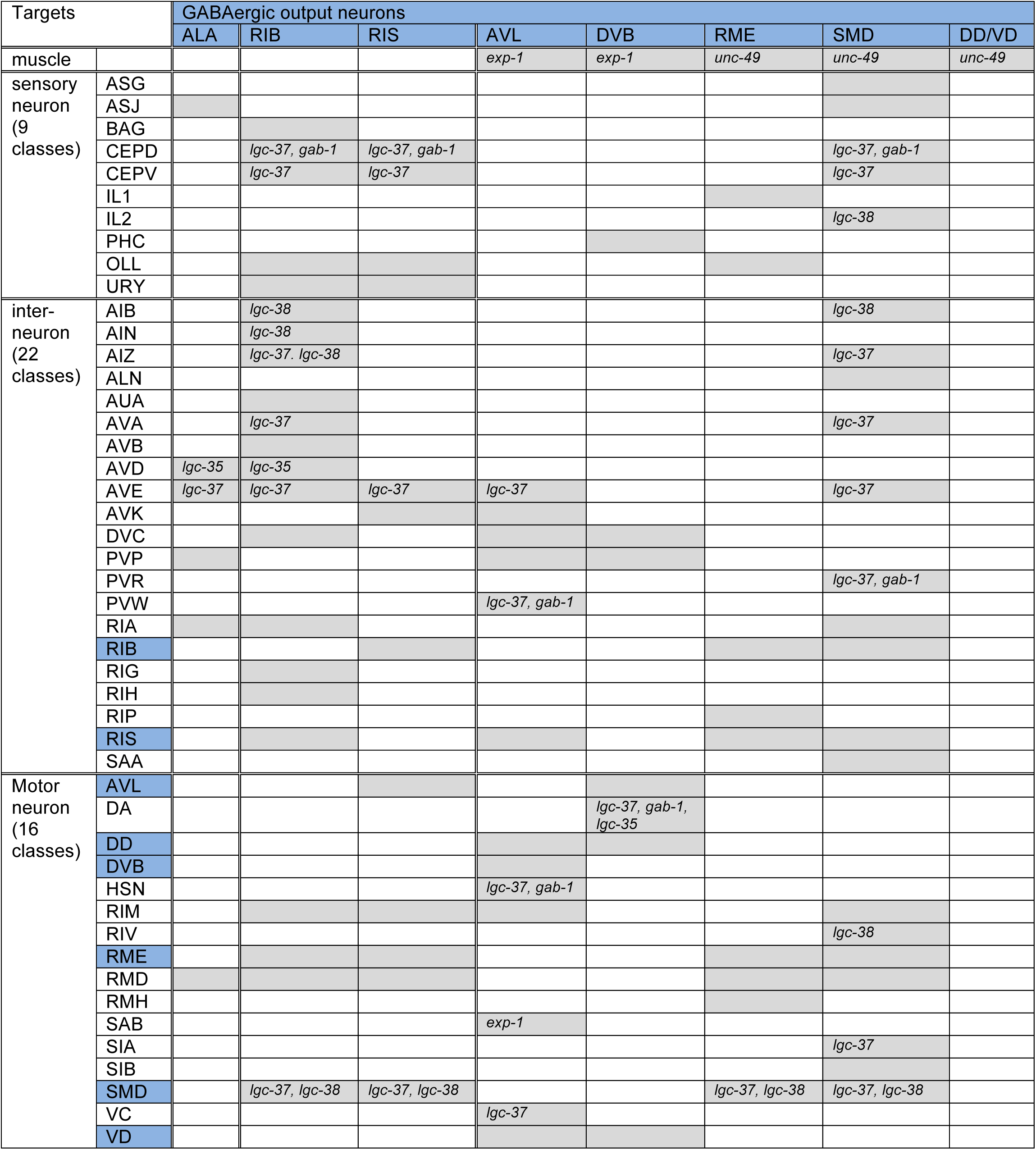
Synaptic targets of GABA(+); UNC-47(+) neurons. Grey box indicates that GABAergic output neuron synapses onto this target cell and genes in the boxes represent expression of the ionotropic GABA_A_-type receptor subunits in the target cell (**Fig.4**). For a complete list of GABA receptor (+) neurons, see **Table 4**. Blue shading indicates that the target neurons is GABA(+). Wiring data is from www.wormwiring.org and includes all synapses observed.

**Table 4:**
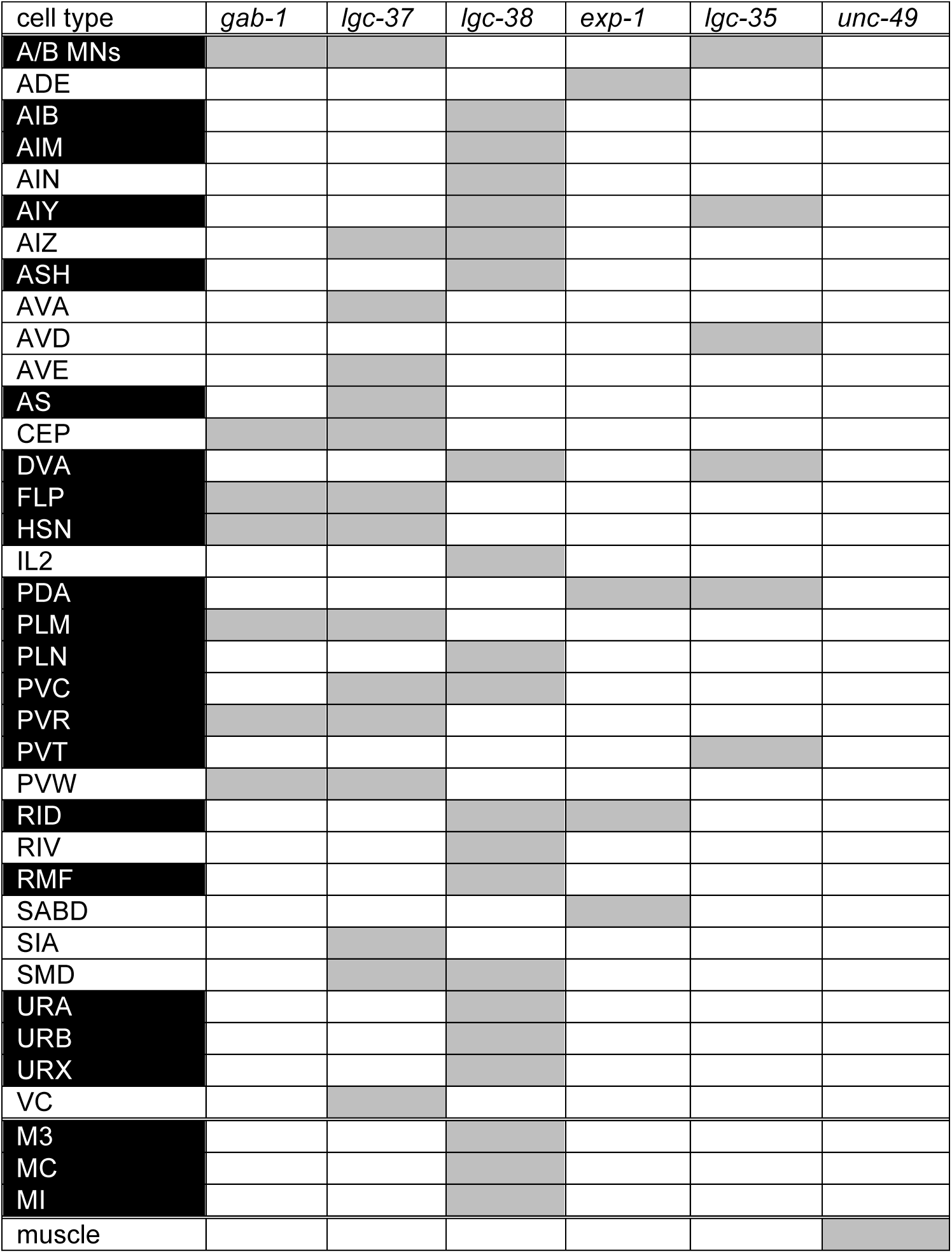
Summary of expression of GABA_A_-type receptors. *gab-1, lgc-37, lgc-38* expression is from this paper (**Fig.4**), *exp-1* is from Beg et al., 2003, *lgc-35* from Jobson et al., 2015 and *unc-49* from Bamber et al., 1999. Cells that are not receiving GABAergic synaptic input (as shown in wormwiring.org) are boxed black and likely receive GABA spillover signals. Pharyngeal neurons and muscle cells are separated by a double line.

Given the relatively small number of GABAergic neurons, it is notable that 28 out of the 49 neurons that receive synaptic input from GABAergic neurons receive such inputs from more than one GABAergic neuron; in many cases inputs are received from more than half of all the distinct GABAergic neuron classes (**Table 3**). For example, the RMD head motor neurons are innervated by the GABAergic RME and SMD motorneurons, by the RIS and RIB interneurons and by the ALA sensory neuron. The GABAergic RIS and RIB interneurons co-innervate a number of distinct neuron classes. Apart from the RMD head motorneurons, the OLL and URY head sensory neurons, the AVE command interneuron and the RIM, RME and SMD head motor neurons are co-innervated by RIS and RIB.

Moreover, all except one of the GABA(+) neurons (the unusual ALA GABA reuptake sensory neurons), receive GABAergic input from other GABAergic neurons (**Table 3**). GABAergic disinhibition, a common organizational principle of inhibitory circuits in the vertebrate CNS (Roberts, 1974) is therefore also observed in the *C. elegans* nervous system.

### GABA receptor expression

The 48 neurons that are postsynaptic to GABA(+); UNC-47(+) neurons are candidates to express at least one of the seven GABA_A_-type receptor encoded by the *C. elegans* genome (Hobert, 2013). Three of these receptors have been previously examined, *unc-49, exp-1* and *lgc-35* (Bamber et al., 1999; Beg and Jorgensen, 2003; Jobson et al., 2015). UNC-49 and EXP-1 are expressed in the muscle cells innervated by the GABAergic motor neurons (Bamber et al., 1999; Beg and Jorgensen, 2003). EXP-1 is also expressed in the SAB neurons, innervated by the GABAergic AVL neurons and LGC-35 is expressed in the AVD neurons, innervated by ALA and RIB (Beg and Jorgensen, 2003; Jobson et al., 2015). However, the expression of the most canonical GABA_A_-type receptors in the worm genome (shown in **Fig.4A**)(Tsang et al., 2007), the two alpha-subunit type GABA_A_ receptors LGC-36 and LGC-37 and the beta-subunit GABA_A_ receptor GAB-1, has not previously been reported. We examined the expression of two of these alpha or beta subunit GABA receptors, *gab-1* and *lgc-37*, and we examined the expression of the *unc-49*-related *lgc-38* gene, using transcriptional reporter gene fusions. All three reporter transgenes are expressed in a restricted number of head and tail neurons (**Fig.4B-D;** summarized in **Table 4**). Overlaps in expression suggest that the respective receptors form heteromeric receptors. We mapped neurons that express any of the GABA_A_-type receptors onto a matrix of neurons innervated by GABA neurons, as shown in **Table 3** (and as indicated by blue letter coding in **Fig.4B-D**). This expression data provides a possible starting point to disrupt GABAergic signaling in a synapse-specific manner.

**Fig.4:**
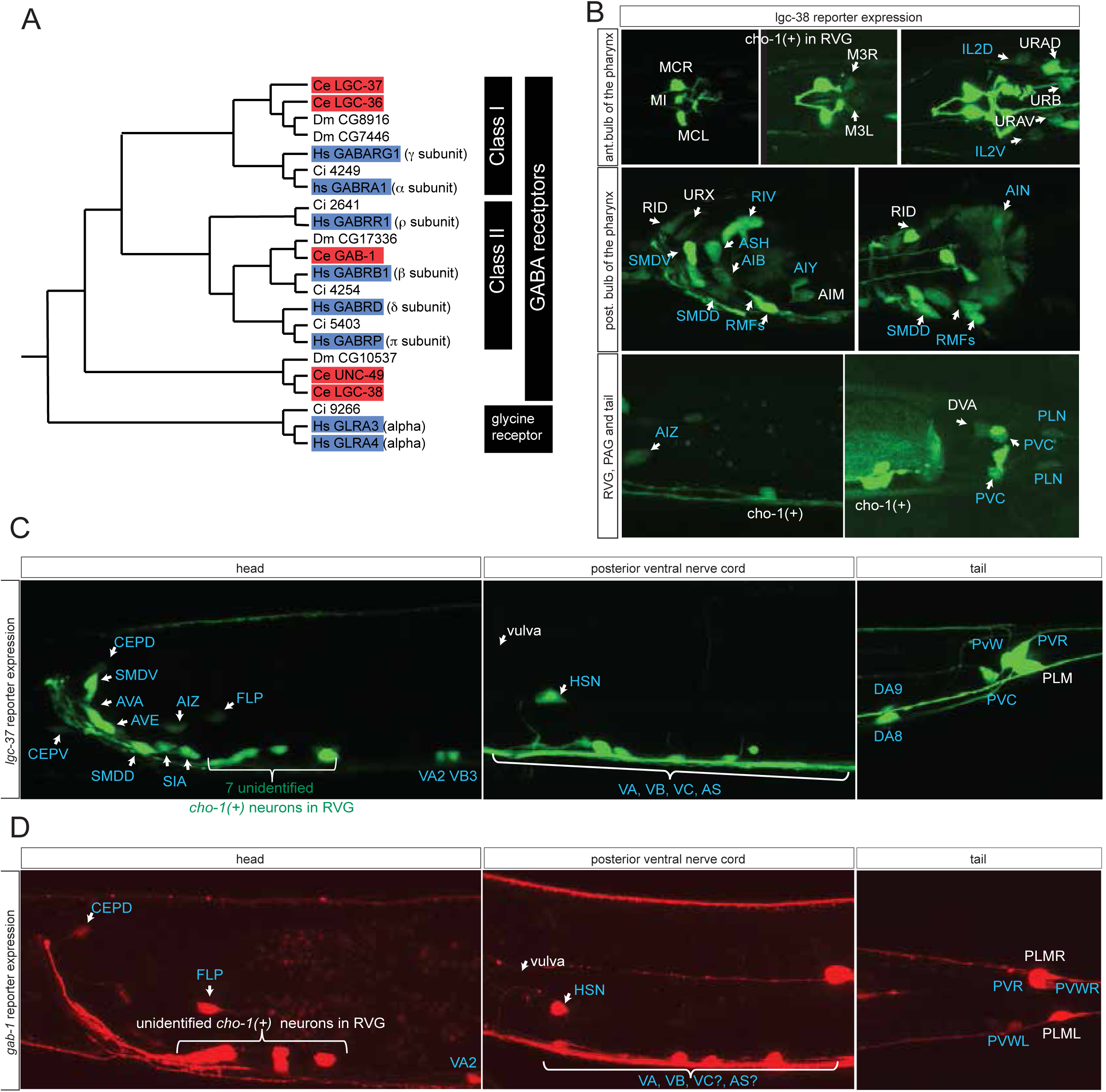
Ionotropic GABA receptor expression. **A:** Phylogeny of the *C. elegans* GABA receptor-encoding genes (red boxes) that are most similar to canonical human GABA receptor-encoding genes (blue boxes). The dendrogram has been adapted from (Tsang et al., 2007). Two other bona-fide GABA receptors, EXP-1 and LGC-35, are more distantly related, falling outside this part of the tree. **B-D:** Expression of *lgc-38* (B) *lgc-37* (C) and *gab-1* (D) reporter constructs. Blue labels indicate neurons innervated by a GABA-positive neuron (see **Table 3**). Cells were identified with *unc-47, eat-4* or *cho-1* reporters as landmarks in the background. These transcriptional reporter gene fusions (see Materials and Methods) do not capture the entire loci and hence may lack regulatory information.

In addition to being expressed in neurons that show anatomic innervation by GABAergic neurons, it is clearly evident that GABA_A_-type receptors are also expressed in neurons that are not anatomically connected to GABA-positive neurons (**Fig.4B-D**, white labels; all neurons listed in **Table 4**). These receptors likely mediate GABA spillover transmission, a phenomenon observed in the vertebrate CNS as well as in *C. elegans* (Jobson et al., 2015; Rossi and Hamann, 1998). Specifically, the previously described expression patterns of *exp-1* and *lgc-35* reveal expression in non-GABA-innervated neurons (*exp-1*: ADE, RID; *lgc-35*: A/B-type MNs, DVA, PVT, AIY). Our expression analysis further extends this notion (**Fig.4B-D;** neurons boxed in black in **Table 4**), identifying expression of, for example, *lgc-37* in the cholinergic A/B-type neurons that were previously reported to receive spill-over GABA signals (Jobson et al., 2015). Spillover transmission may even extend into the pharyngeal nervous system, where we detect GABA_A_-type receptor expression (**Table 4**), but no GABA staining.

### The Tailless/Tlx orphan nuclear receptor NHR-67 controls the GABAergic identity of a diverse set of GABAergic neurons

We used the expanded map of the GABAergic nervous system as a starting point to elucidate transcriptional regulatory programs that specify GABAergic neuron identity. Previously, the effect of a number of transcription factors on GABAergic identity has been described, but in most cases, the analysis has either been limited to a few markers or a number of important questions about the specificity of the involved factors has remained unanswered. We addressed the functions of these factors systematically in this and the ensuing sections.

Sarin et al. have previously shown that the Tailless/Tlx orphan nuclear receptor *nhr-67* is expressed in the GABAergic RMEs, RIS and AVL neurons and that *nhr-67* expression is maintained in these neurons throughout adulthood in the RME and RIS neurons (Sarin et al., 2009). Consistent with a possible role of *nhr-67* as a regulator of GABAergic identity, it was also reported that loss of *nhr-67* affected expression of *unc-47/VGAT* in RME, AVL and RIS (Sarin et al., 2009). However, whether *nhr-67* affects GABA synthesis, or the expression of other GABAergic identity features was not tested. We first isolated an unambiguous molecular null allele of *nhr-67, ot795*, using Mos transposon-mediated gene deletion (MosDEL; see Experimental Procedures)(**Fig.5A**). *nhr-67(ot795*) animals display an embryonic/L1 arrest phenotype. We stained these null mutant animals for GABA and examined *unc-25/GAD*, *unc-47/VGAT* and *unc-46/LAMP* expression in these mutants. We found abnormalities in the expression of all markers in all three, normally *nhr-67-*expressing neuron classes, AVL, RME and RIS (**Fig.5B**). In some cases, defects are modest, but as we will describe in the next section, are strongly enhanced by removal of factors that likely cooperate with *nhr-67*.

**Fig.5:**
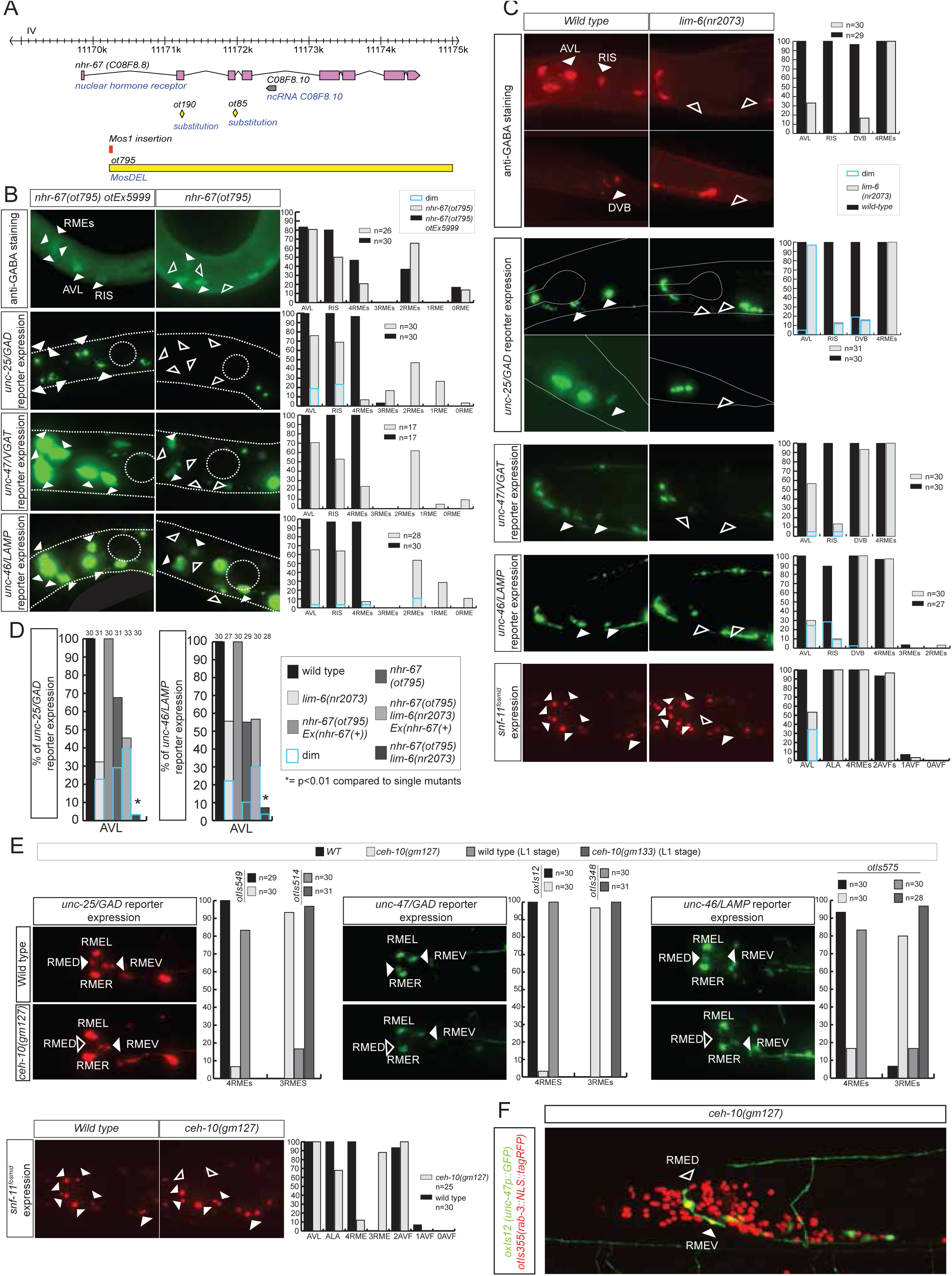
*nhr-67* cooperates with distinct homeodomain proteins in distinct GABAergic neuron types. **A:** *nhr-67* locus with new null allele. **B:** Expression of GABA pathway genes and anti-GABA staining of *nhr-67* null mutant animals that contain or do not contain a rescuing array (*otEx5999*) which contains a wild-type copy of the *nhr-67* locus. **C:** *lim-6* null mutant animals display defects in GABAergic neuron identity. **D:** *nhr-67* and *lim-6* genetically interact in AVL. **E:** *ceh-10* mutant animals display defects in GABAergic neuron identity. Null mutants (*gm133)* were scored as L1, when they arrest. Viable *gm127* hypomorphic mutants were scored as adults. Wildtype data in panel C is re-iterated in panel E. **F:** *ceh-10* does not affect the generation of RMED as assessed by detection of RMED with a pan-neuronal marker.

### *nhr-67* cooperates with distinct homeobox genes in distinct GABAergic neurons

Two previous studies had implicated two homeobox genes in the regulation of GABA identity of a subset of the *nhr-67* expressing neurons, the Prd-type homeobox gene *ceh-10* and the LIM homeobox gene *lim-6. ceh-10* was found to affect GABA staining and *unc-25/GAD* expression in the RMED neurons (where *ceh-10* was reported to be expressed)(Forrester et al., 1998; Huang et al., 2004). We extended this previous finding by demonstrating that in *ceh-10* mutants, not only the *unc-25* reporter, but also the *unc-47* and *snf-11* reporters fail to be expressed in RMED (**Fig.5E**). Absence of expression of these reporter is not reflective of lineage defects, since we find that in *ceh-10* hypomorphic mutants in which GABA staining is also severely affected, the RMED neurons are nevertheless formed, as assessed by examination of a pan-neuronal marker (**Fig.5F**). In addition to affecting expression of GABAergic markers, *ceh-10* also affects the expression of an *nhr-67* fosmid based reporter (**Fig.5 – supplement 1**).

The LIM homeobox gene *lim-6* was found to control *unc-25* expression in the RIS, AVL and DVB neurons (Hobert et al., 1999). We corroborated the impact of *lim-6* on GABA identity by anti-GABA staining of *lim-6* null mutant animals (which had not previously been done), finding that GABA staining is affected in AVL, RIS and DVB (**Fig.5C**). Since *nhr-67* null mutants did not display fully penetrant defects in the AVL and RIS neurons, we tested whether *nhr-67* and *lim-6* may collaborate in the specification of these neurons. We find that in *nhr-67; lim-6* double null mutants, the AVL neurons show synergistic defects in GABAergic identity specification (**Fig.5B**). *lim-6* expression is not affected in AVL and RIS of *nhr-67* mutants, but *lim-6* (as well as *ceh-*10) controls *nhr-67* expression (**Fig.5 – supplement 1**). Taken together, *nhr-67* appears to collaborate with distinct homeobox genes in distinct neurons and appears to be regulated by these factors. Since both *ceh-10* and *lim-6* remain expressed throughout the life of the respective neurons, we surmise that *ceh-10* and *lim-6* induce a critical cofactor (*nhr-67*) that they then work together with. In other cases of collaborating transcription factors, it has also been demonstrated that one factor acts upstream of the other to then cooperate with the induced factor (e.g. *ttx-3* induces *ceh-10* expression and both factors then cooperate to drive cholinergic identity of the AIY neurons; (Bertrand and Hobert, 2009; Wenick and Hobert, 2004)).

### The homeobox gene *tab-1* controls GABAergic identity of the RMEL/R neurons

While *lim-6* and *ceh-10* homeobox genes may work together with *nhr-67* in subsets of GABAergic neurons to control GABAergic identity, no cooperating factor for *nhr-67* function in the RMEL/R neurons was apparent. We screened several dozen homeobox mutants for defects in GABA staining without success (data not shown) and then screened for EMS-induced mutants in which *unc-47* expression in the RME neurons was absent (see Methods). We identified a mutant allele, *ot796*, in which *unc-47* failed to be expressed in the left and right RME neurons. Whole genome sequencing revealed that *ot796* contains a splice site mutation in the *tab-1* locus (**Fig.6; Fig.6 – supplement 1**; **Table 5**). The *ot796* mutant allele fails to complement the GABA differentiation defects of other mutant alleles of *tab-1* and three additional alleles of *tab-1* display similar *unc-47* expression defects as *ot796* (**Table 5**). *tab-1* mutants also fail to properly express *unc-25* and *unc-46* in RMEL/R and show loss of anti-GABA staining. A *tab-1* fosmid-based reporter is expressed in both the left and right RME neurons (**Fig.6**).

**Fig.6:**
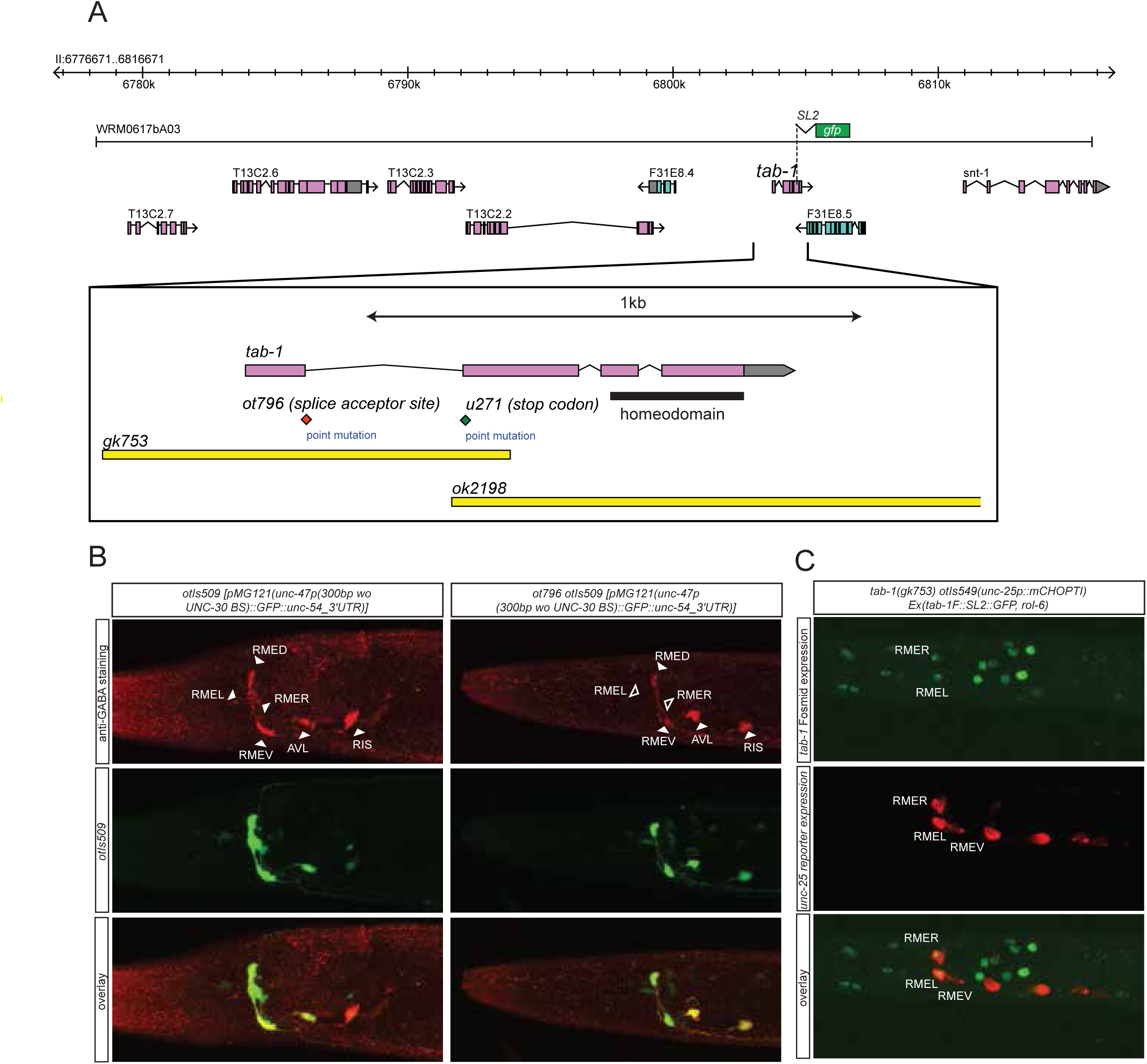
*tab-1* is another homeodomain protein interacting with nhr-67 to specify GABAergic identity. **A:** *tab-1* locus, alleles and fosmid reporter. **B:** *tab-1* affects *unc-47* reporter expression and anti-GABA staining. The *unc-47* reporter used has a presumptive *unc-30* binding site deleted, which diminishes expression of this reporter in VNC D-type neurons. Effects on these and other reporters is quantified in **Table 5**. **C:** Expression pattern of the *tab-1* fosmid reporter indicated in panel A.

**Table 5:**
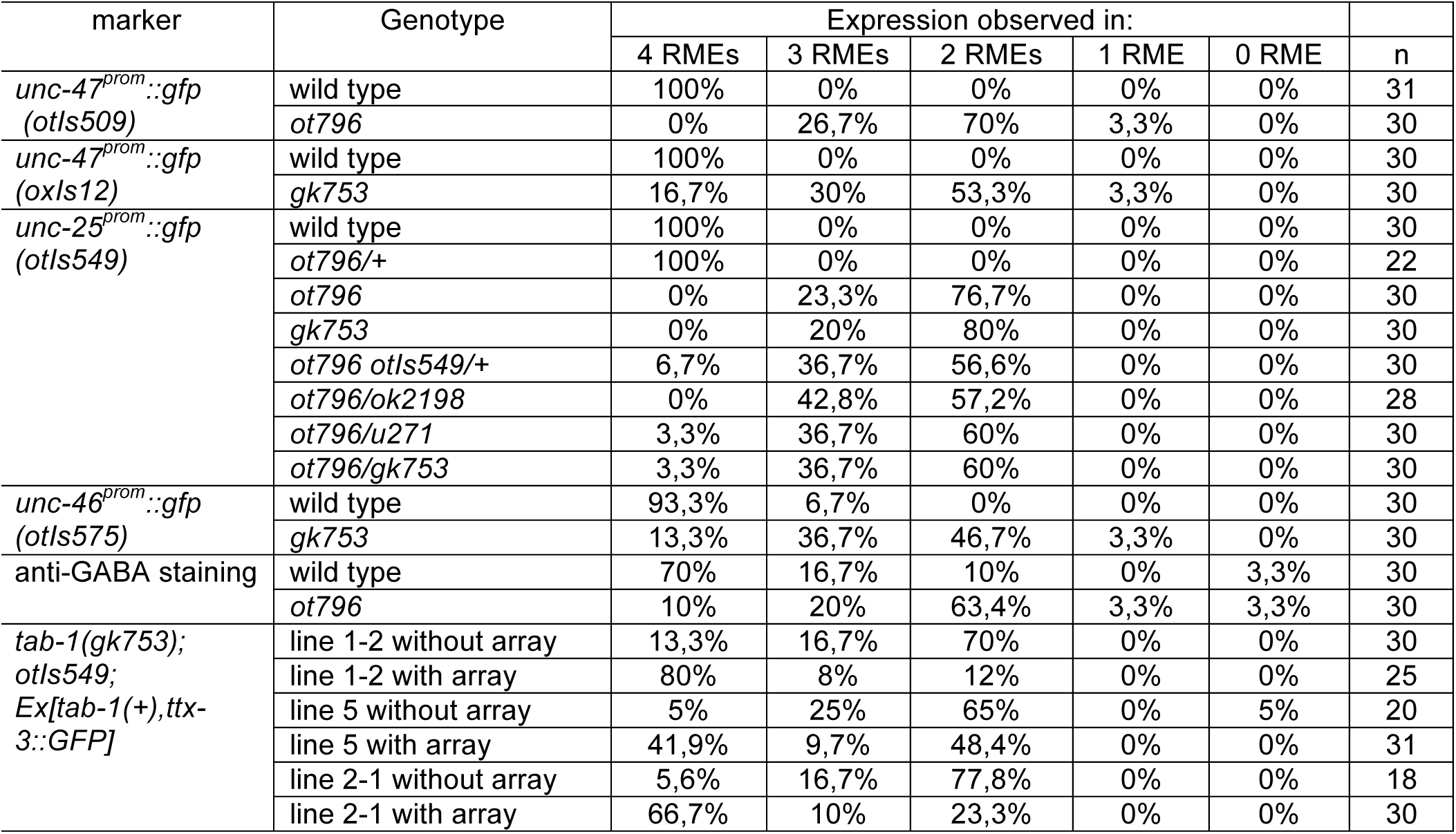
Genetic characterization of *tab-1* function in the RME neurons. Expression in RME can not always been unambiguously assigned to the dorsal, ventral, left or right RME neuron, hence only the total number of RME neurons was counted.

*tab-1* encodes the sole *C. elegans* ortholog of the *Drosophila* Bsh homeobox gene and the vertebrate Bsx genes (Pang and Martindale, 2008). *tab-1* mutants (for “<u>t</u>ouch <u>ab</u>normal”) were previously isolated based on their defects in the touch response (L. Carnell, B. Harfe, A. Fire and M. Chalfie, pers. comm.). *Drosophila* Bsh has been implicated in the specification of several neuron types in the *Drosophila* optic lobe (Hasegawa et al., 2013). A subset of these neurons are GABAergic (Raghu et al., 2013), indicating that a function of Bsh-type homeobox genes in the specification of the GABAergic phenotype may be phylogenetically conserved.

### The GATA2/3 ortholog *elt-1* affects GABA identity of D-type motorneurons

To identify additional regulators of GABAergic identity, we systematically examined the function of *C. elegans* orthologs of genes known to regulate GABAergic identity in the CNS of vertebrates. We examined possible functions of such *C. elegans* orthologs by examining their expression and mutant phenotypes and found a striking example of conserved function. The vertebrate GATA2 and GATA3 transcription factors operate as selector genes of GABAergic identity in several distinct regions of the vertebrate CNS, including the spinal cord, midbrain, forebrain and hindbrain (Achim et al., 2014; Joshi et al., 2009; Kala et al., 2009; Lahti et al., 2016; Yang et al., 2010). GATA2 may act transiently after the generation of GABAergic neurons and then pass on its function to the GATA3 paralog. The sole *C. elegans* ortholog of vertebrate GATA2/3 is the *elt-1* gene (Gillis et al., 2008), which controls early hypodermal fate patterning (Page et al., 1997). We found that an *elt-1* fosmid-based reporter gene is expressed in all D-type motor neurons throughout their lifetime, but not in any other GABA-positive neuron (**Fig.8A**). Since *elt-1* mutants display early embryonic lethality, we conducted genetic mosaic analysis to examine the effect of loss of *elt-1* function on D-type motor neurons, some of which generated only postembryonically (the VD MNs). We balanced *elt-1* null mutants with a fosmid that contains the *elt-1* locus and an *unc-47::gfp* reporter to assess the loss of this marker in GABAergic D-type neurons. We found that live animals that lost the rescuing array show defects in *unc-25/GAD, unc-47/VGAT* and *unc-46/LAMP* expression in D-type neurons (**Fig.8B**). The loss of expression of GABA markers is not a reflection of loss of the cells, since expression of *unc-30*, the presumptive regulatory co-factor for *elt-1* is still normally express in the D-type neurons of *elt-1* mutants (**Fig.8C**).

**Fig.8:**
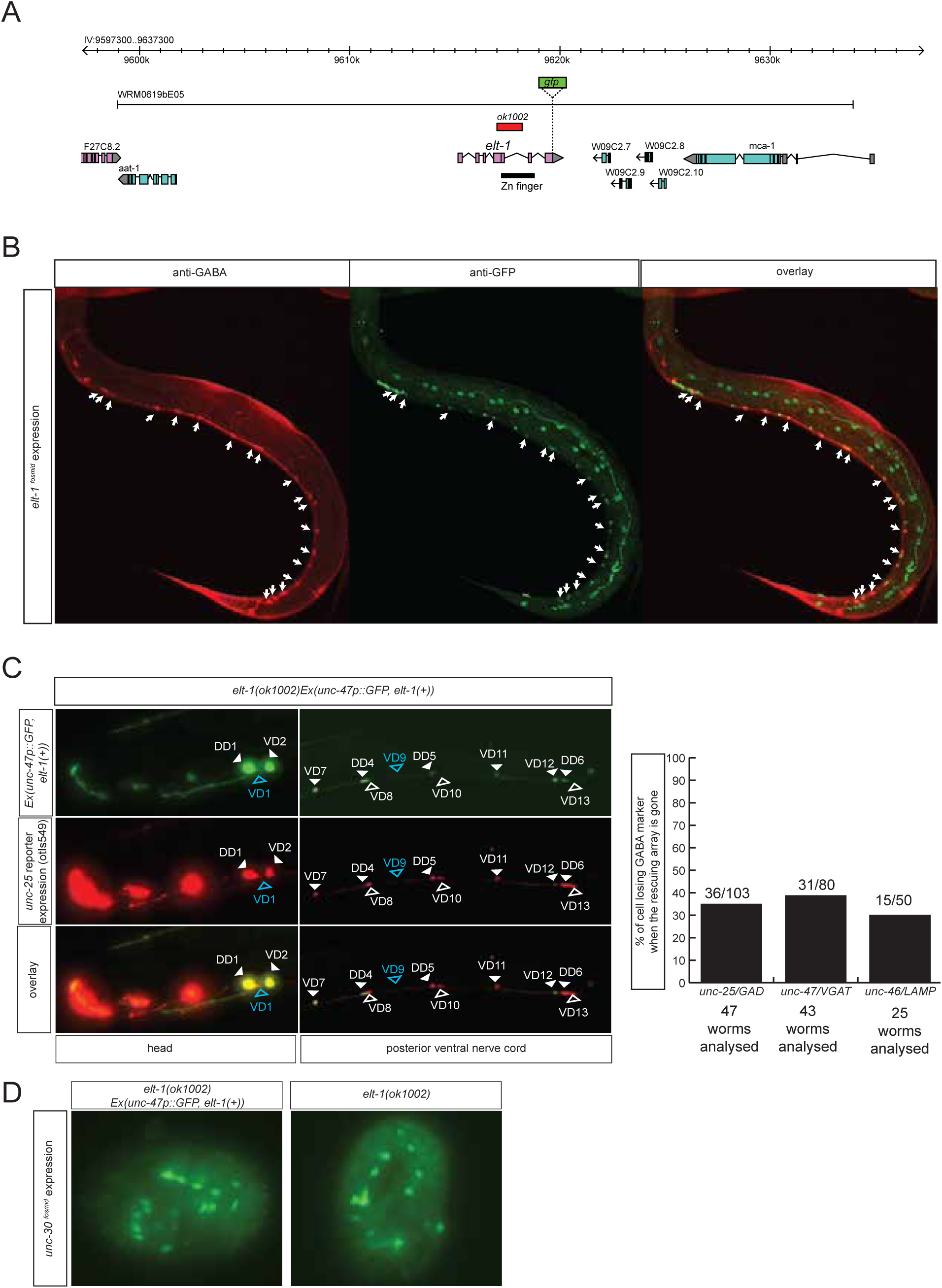
The GATA2/3 ortholog *elt-1* controls GABAergic identity of D-type MNs. **A:** *elt-1* locus and fosmid reporter used for expression analysis. The same, but untagged fosmid was used for mosaic analysis (panel C). **B:** Expression pattern of the *elt-1* fosmid based reporter construct. **C:** Mosaic analysis of *elt-1* function. *elt-1* null mutants that carry a rescuing array, as well as an array markers that labels the presence of the array in GABA neurons (*unc-47::gfp*) is scored for *unc-25::mChOPTI* expression. Whenever no *gfp* signal is observed in a D-type motorneuron, the expression of *unc-25::rfp* is scored (bar graph). **D:** *unc-30::gfp* expression is not affected in *elt-1* mutants.

Two other vertebrate genes controlling GABAergic identity in the vertebrate CNS (Tal1/2 and Lbx) have no *C. elegans* ortholog. The *C. elegans* orthologs of two prominent other regulators of GABAergic neuron identity in vertebrates, the Dlx genes (*ceh-43* in *C. elegans*), and Ptf1a (*hlh-13* in *C. elegans*) have no role in GABAergic identity control since *ceh-43* is not expressed in mature GABAergic neurons (data not shown) and since *hlh-13* null mutants show no defects in GABA staining (data not shown).

### Homeobox genes controlling the identity of GABA reuptake neurons

We next sought to identify factors that control the identity of the GABAergic reuptake neurons that we newly identified. The *ceh-14* LIM homeobox gene and the *ceh-17* Prd-type homeobox gene were previously shown to cooperate in the specification of several, peptidergic terminal identity features of the GABA reuptake neuron ALA (Van Buskirk and Sternberg, 2010). We find that GABA staining of the ALA neurons is abrogated in *ceh-14* and *ceh-17* mutants (**Fig.7A**). Since lack of GABA staining is expected to be due to the failure to express *snf-11* we crossed the *snf-11* fosmid reporter into *ceh-14* mutants and found its expression to be abrogated in the ALA neuron (**Fig.7A**).

**Fig.7:**
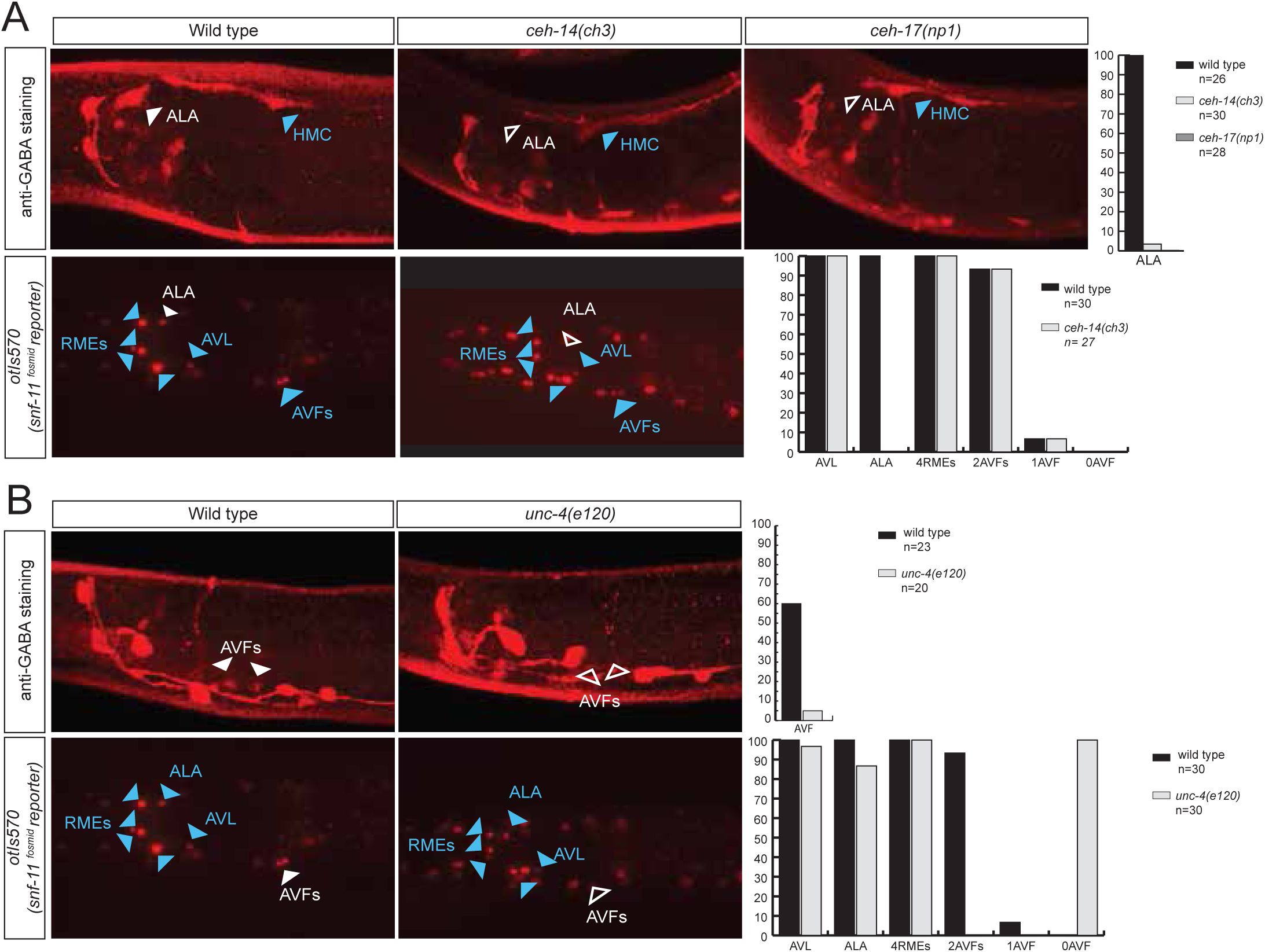
Homeobox genes controlling the identity of GABA reuptake neurons. **A:** *ceh-14* and *ceh-17* specify the identity of the newly identified ALA GABAergic neuron, as assessed by GABA staining (upper panels) and *snf-11* reporter gene expression (lower panels) **B:** *unc-4* affects GABA staining and *snf-11* reporter gene expression in the AVF neuron. unc-4 affects GABA, snf-11

The AVF reuptake neuron was previously shown to express the Prd-type homeobox gene *unc-4* (Miller and Niemeyer, 1995). We find that *unc-4* mutants do not show GABA staining in AVF and, as a likely reason for the absence of GABA staining, fail to express the *snf-11/GAT* transporter (**Fig.7B**). *unc-3*, a transcription factor controlling the identity of P-cell derived cholinergic neurons, is also expressed in the AVF neurons (derived from P and W), but does not affect anti-GABA staining in AVF (data not shown).

### GABAergic neurotransmitter identity is coupled with the adoption of other identity features

Lastly, we set out to address the question whether factors that control GABAergic identity are committed to only control GABAergic identity or whether they also control additional identity features of the respective GABAergic neurons. In other words, is GABAergic neurotransmitter coupled to the adoption of other identity features? This appears to indeed be the case if one considers previously published results. Specifically, *ceh-14* and *ceh-17* were previously described to control several identity features of ALA (Van Buskirk and Sternberg, 2010) and, as mentioned above, we show here that it also controls GABA identity (**Fig.7**). Similarly, *lim-6*, which specifies GABA identity of RIS and AVL, had previously been found to control several identity aspects of the RIS interneuron, namely expression of two biogenic amine receptors, a glutamate receptor and an Ig domain protein (Tsalik et al., 2003). We found that *lim-6* also controls the expression of a more recently identified RIS marker, *nlr-1* and *lim-6* affects *flp-22* expression in AVL (**Fig.9A**). Together with the GABA staining defects of RIS and AVL in *lim-6* mutants described here, this demonstrates that *lim-6* coregulates GABA identity acquisition and other identity features.

**Fig.9:**
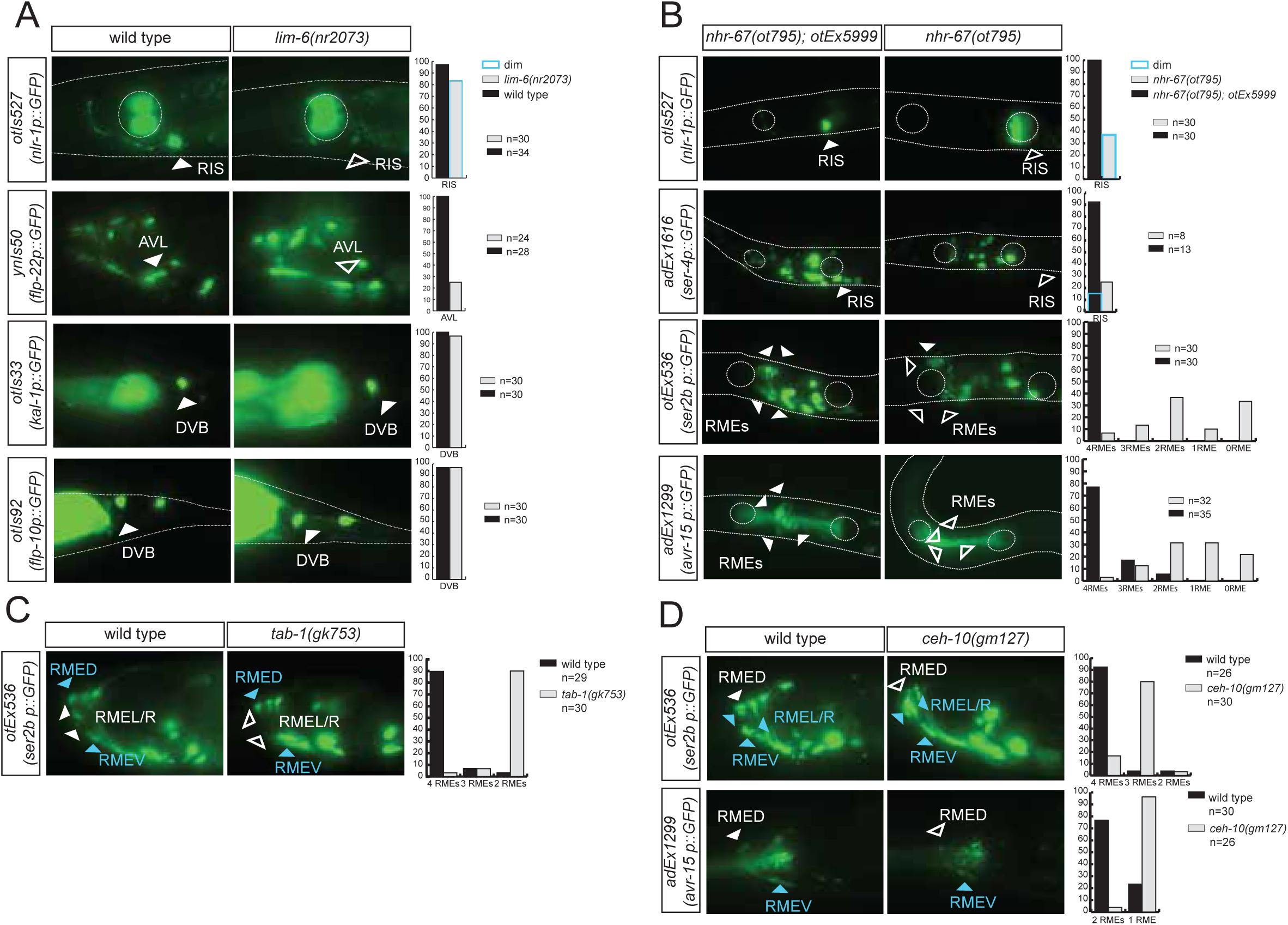
Transcription factors controlling GABAergic identity also control other cellular identity markers. **A:** *lim-6* controls cell identity markers in AVL and RIS, but not DVB. **B:** *nhr-67* controls additional neuron identity markers of GABAergic neurons. *nhr-67(ot795)* animals are not viable and *otEx5999* is an extrachromosomal array that contains copies of the wild-type *nhr-67* locus as well as an array marker (*unc-47prom::mCHOPTI*). Animals that either contain this array or do not contain it were scored for mutant phenotypes at the L1 stage. **C:** *tab-1* controls RMEL/R identity markers. **D:** *ceh-10* controls RMED identity markers.

We corroborated the notion of coregulation further by examining whether *nhr-67* not only controls GABA identity but also other identity features. We find that in the RIS neurons, *nhr-67* also controls the expression of the 5HT receptor *ser-4* and of *nlr-1* (**Fig.9B**). In the RME neurons, *nhr-67* affects not only GABA identity, but also expression of the tyramine receptor *ser-2* and the Glu-gated ion channel *avr-15* (**Fig.9B**). *ser-2* and *avr-15* expression is also affected in RMED by *ceh-10* and *ser-2* expression in RMEL/R is affected by *tab-1* (**Fig.9C,D**).

One notable exception to the coregulatory theme appears in the DVB motorneurons. GABA staining is absent in *lim-6* mutants (**Fig.5C**) and *unc-25/GAD* expression is also severely affected (**Fig.5C**)(Hobert et al., 1999). However, neither *unc-47::gfp* expression is affected in *lim-6* null mutants, nor the expression of two additional DVB markers, *kal-1* and *flp-10* (**Fig.9**). We note that in the AVL neuron, the effect of *lim-6* on some markers is also very modest, but significantly enhanced if the combinatorial cofactor for *lim-6* in AVL, *nhr-67*, is also removed (**Fig.5**). We therefore suspect that *lim-6* may act in a partially redundant manner with a cofactor in DVB as well.

In conclusion, in most cases examined, regulatory factors that control GABAergic identity features also control other identity features of the neurons examined.

### The *unc-42* homeobox gene represses GABA identity in several types of motor-and interneurons

In our search for additional regulators of the GABAergic phenotype we noted that the normally peptidergic AVK neurons ectopically stain with anti-GABA antibody in animals that lack the *unc-42* homeobox gene (**Fig.10**). Ectopic *unc-25/GAD, unc-46/LAMP* and *snf-11/GAT* reporter gene expression is also observed in the AVK neurons of *unc-42* mutants (**Fig.10**). Previous work had shown that in *unc-42* mutants several markers of AVK fail to be expressed (Wightman et al., 2005). It therefore appears that *unc-42* promotes the peptidergic identity of AVK and suppresses an alternative GABAergic differentiation program.

**Fig.10:**
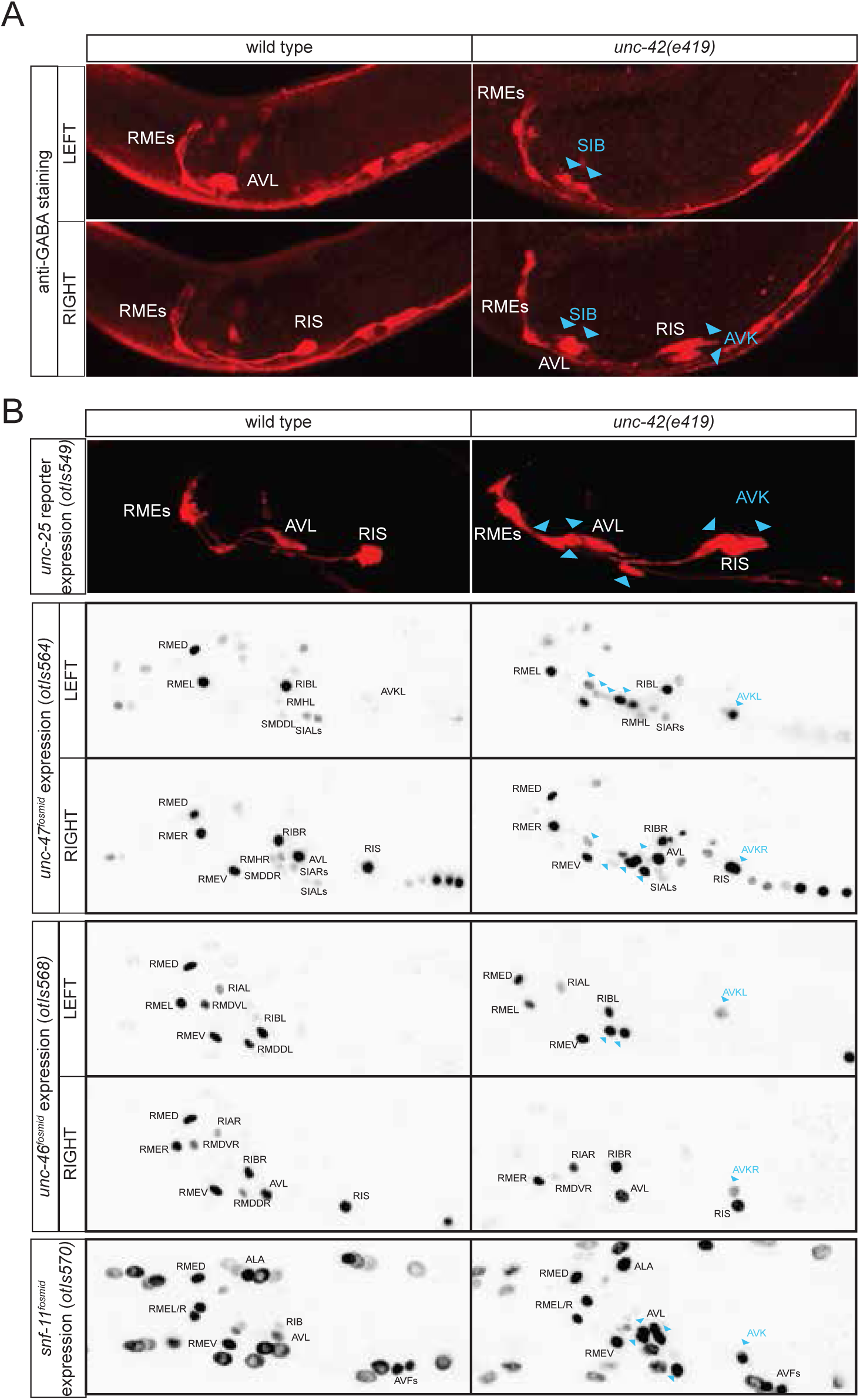
*unc-42* represses GABAergic neuron identity. **A:** Anti-GABA staining in head neurons of wild-type and *unc-42* mutant animals. **B:** Reporter gene expression in head neurons of wild-type and *unc-42* mutant animals. Blue arrows point to ectopic expression.

*unc-42* is also expressed in a cluster of cholinergic motor neurons in the ventral ganglion, posterior to the nerve ring, including the SMDD/V and SIBD/V motor neuron pairs (Baran et al., 1999; Pereira et al., 2015). We observed ectopic GABA staining, as well as *unc-25* and *snf-11* reporter gene expression in this region in *unc-42* mutants (**Fig.10**), indicating that *unc-42* may suppress GABAergic differentiation programs in cholinergic motor neurons as well; the best candidates for the neurons that convert from cholinergic to GABAergic are the normally *unc-42*-expressing SIBD and SIBV motor neuron pairs, based on position (**Fig.10**). Taken together, the existence of a regulatory factor that suppresses GABAergic identity in several distinct neuron types suggests that GABAergic identity was more broadly expressed in an ancestral nervous system, but suppressed by the recruitment of a factor that could impose an alternative identity on these neurons.

## DISCUSSION

### A *C. elegans* neurotransmitter atlas

This is the third and last paper in a trilogy of mapping papers that chart the three main neurotransmitter systems in *C. elegans*, Glu, ACh and GABA (Pereira et al., 2015; Serrano-Saiz et al., 2013). The maps of the major three fast transmitter system constitute an atlas of neurotransmitter usage whose breadth is unprecedented in any other nervous system. The atlas is shown in **Fig.11A**, all neurons are listed in **Table 6** and a 3D rendering of this atlas is shown in **Movie 1**. In total, a neurotransmitter identity has now been assigned to 104 out of the 118 neuron classes of the worm. 98 of these employ a “classic” fast neurotransmitter (Glu, GABA, ACh), 6 employ exclusively a monoaminergic transmitter. Several of the neurons using a fast-acting transmitter also cotransmit a monoamine. The 14 neuron classes for which no classic neurotransmitter system has been identified yet may be dedicated to the use of neuropeptides.

**Fig.11:**
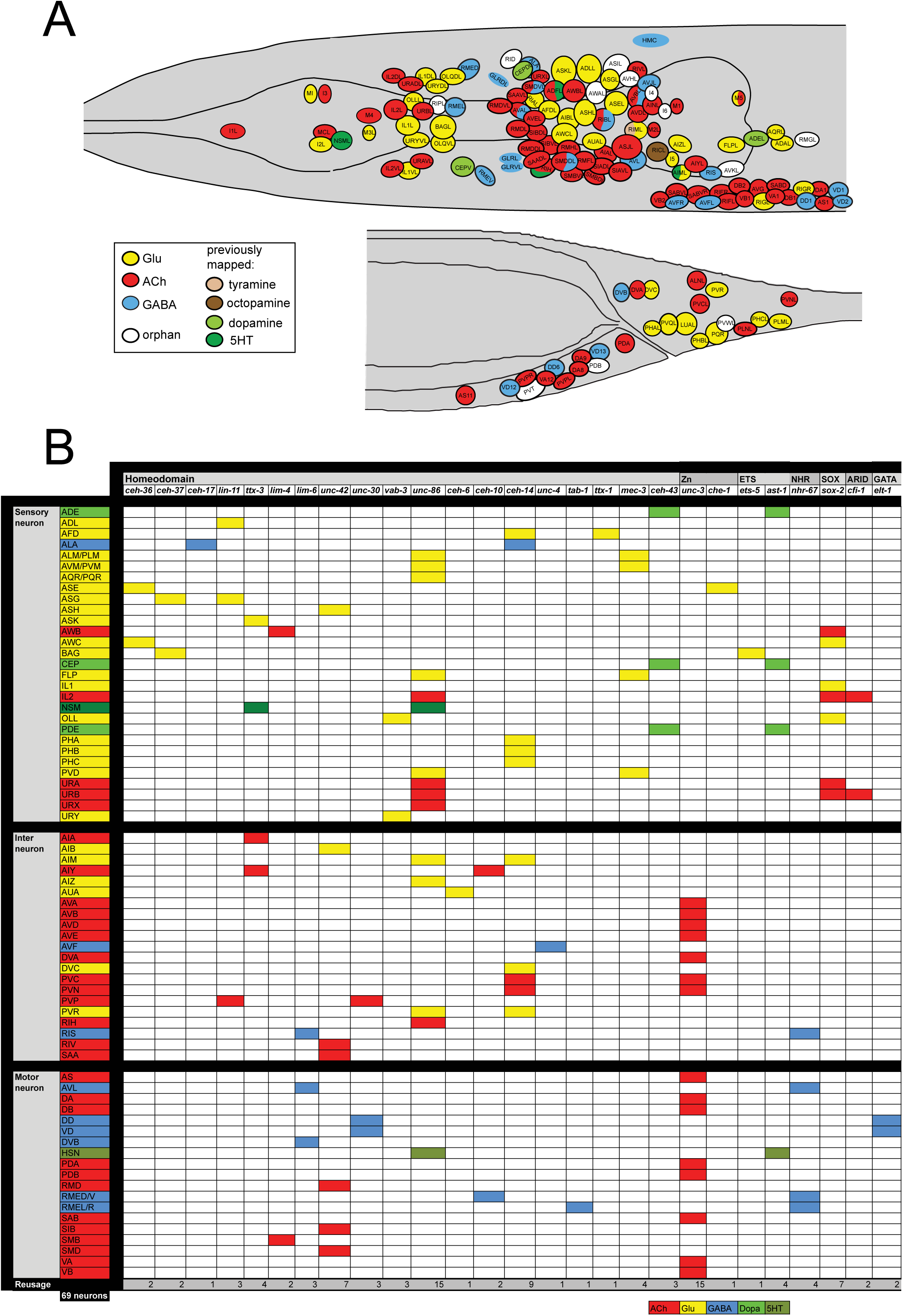
Neurotransmitter maps. **A:** Current status of the *C. elegans* neurotransmitter atlas. Only the head and tail of the worm are shown. The Glu and ACh maps come from (Pereira et al., 2015; Serrano-Saiz et al., 2013) and have been updated here with the GABA neuron analysis. See **Table 6** for complete list. **B:** Regulatory map of neurotransmitter specification. Data is from this paper for GABA and from following papers for additional neurons: (Alqadah et al., 2015; Altun-Gultekin et al., 2001; Doitsidou et al., 2013; Flames and Hobert, 2009; Kratsios et al., 2015; Kratsios et al., 2011; Pereira et al., 2015; Serrano-Saiz et al., 2013; Sze et al., 2002; Vidal et al., 2015; Zhang et al., 2014)

**Table 6:**
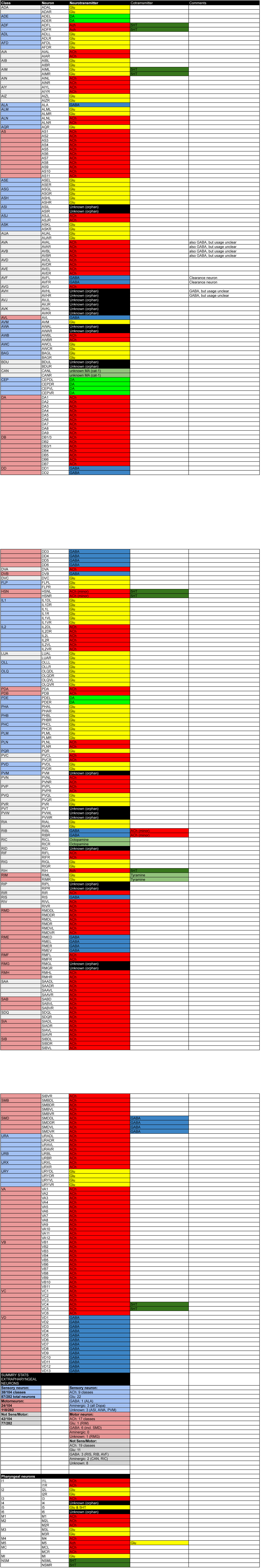
List of all neurotransmitters

### Usage of GABA throughout the nervous system

Among the most notable aspects of this atlas is the previously discussed broad usage of ACh, employed by 52 sensory, inter- and motorneuron classes (out of a total of 118 classes)(Pereira et al., 2015) and the apparent paucity of GABA usage. Only 10 neuron classes use GABA for synaptic signaling (based on expression of the vesicular transporter) and of those, only two are pure interneurons (RIS and RIB ring interneurons; since AVB, AVD and AVF express no known release machinery, we do not consider them as conventional GABAergic interneurons). Similarly, in the complex wiring diagram of the 81 male-specific tail neurons (Jarrell et al., 2012), GABA is used sparsely, namely, by the 4 male-specific EF interneurons, as well as 3 ray sensory neuron pairs.

The paucity of GABA usage contrasts the much broader usage of GABA in the vertebrate CNS, in which 30-40% of all synapses contain GABA (Docherty et al., 1985). However, the apparent paucity of GABA usage in *C. elegans* is no reflection of paucity of inhibitory neurotransmission in *C. elegans*. First, because of the existence of ACh and Glu-gated chloride channels (Dent et al., 2000; Hobert, 2013; Putrenko et al., 2005), GABA is not the only inhibitory neurotransmitter in *C. elegans*. Second, while only relatively few neurons are GABA positive, a much larger number of neurons may be responsive to GABA. This can be inferred from the expression patterns of ionotropic GABA_A_-type neurotransmitter receptors, which extends beyond the limited number of neurons that are innervated by GABA-positive neurons. Such expression supports extensive spill-over transmission, beyond what has previously been observed in the ventral nerve cord (Jobson et al., 2015). Another indication for the breadth of spill-over transmission is the notable restriction of expression of the GABA reuptake transporter GAT. In vertebrate, these transporters are expressed widely throughout the CNS, with most GABA-positive neurons also expressing GAT (GAT1 or GAT3); many postsynaptic targets of vertebrate GABAergic neurons also express GAT (Conti et al., 2004; Swan et al., 1994; Zhou and Danbolt, 2013). In notable contrast, the sole *C. elegans* GAT ortholog SNF-11 is only expressed in a small fraction of the GABA-positive neurons and it is only expressed in two types of GABAergic targets cells (AVL as a target of DVB and body wall muscle as targets of D-type motorneurons). We hypothesize that the restricted expression of SNF-11 is a reflection of GABA not being immediately cleared after release, but is being able to spill over to inhibit non-synaptic targets.

Some of the GABAergic neurons that we newly identified here cotransmit ACh (one of them, the RIB interneuron only expresses very low level of the ACh synthesizing and transporting machinery)(Pereira et al., 2015). ACh/GABA-cotransmitting neurons have been observed in multiple neuron types of the vertebrate CNS as well (Granger et al., 2016). Neurons that use two neurotransmitters can, in principle, package both neurotransmitters into the same vesicles or they can be packaged separately into spatially segregated presynaptic zones (Vaaga et al., 2014). The GABA-and ACh-positive SMD neurons synapse onto two fundamentally distinct cell types – head muscles and a number of distinct inter- and motorneurons (**Table 1**)(White et al., 1986) and may differentially segregate ACh and GABA to distinct these target synapses.

### GABA reuptake: GABA recycling and GABA clearance neurons

Our studies define not only novel GABA-synthesizing and GABA-releasing neurons but also neurons that we term “GABA reuptake neurons”. The GABA-positive nature of these neurons critically depends on the GABA reuptake transporter SNF-11/GAT that is expressed in these neurons. Based on the expression of the vesicular GABA transporter UNC-47/VGAT, we propose that GABA reuptake neurons fall into two categories “GABA recycling neurons” and “GABA clearance neurons”. GABA clearance neurons (the AVF neurons) reuptake GABA but because these cells do not express the GABA vesicular transporter UNC-47, they do not appear to be capable of re-utilizing GABA, i.e. packaging GABA in synaptic vesicles for re-release. The axon of AVF extends through the nerve ring and along the ventral nerve cord and AVF may therefore clear GABA released from several different GABAergic neuron types. GABA clearance by AVF may control communication between GABA-releasing neurons and their postsynaptic, GABA receptor-expressing targets. AVF may also restrict and spatially define non-synaptic GABA spillover transmission. Whether analogous GABA clearance neurons exist in the vertebrate CNS is as yet unclear, but it is notable that the vertebrate CNS does contain neurons that do not synthesize GABA but take it up via GAT (Conti et al., 2004; Swan et al., 1994). However, it is generally assumed that these neurons are postsynaptic to GABAergic neurons and hence, that GABA reuptake occurs at the synapse. In contrast, AVF is not a synaptic target of GABAergic neurons.

Another potential type of GABA reuptake neurons not only expresses the GABA reuptake transporter GAT, but also expresses the UNC-47/VGAT vesicular transporter. We speculate that these neurons are possible “GABA recycling neurons” that synaptically release GABA after reuptake. The ALA neuron class falls into this category. ALA, which extends two processes into the nerve ring, may take up GABA released from any of the GABA-releasing neurons that also extend processes into the nerve ring (**Fig.1B**), with SMD neurons being the best candidates due to the direct adjacency of their processes. While we do not have direct evidence that ALA then re-releases GABA, ALA has previously been shown to inhibit the activity of the synaptically connected AVE command interneurons to control locomotory behavior (Fry et al., 2014). Since ALA does not express any other known fast transmitter system, we posit that this inhibitory activity is mediated by GABA released by ALA and perceived by GAB-1, an ionotropic GABA receptor we find to be expressed in AVE. GABA reuptake by ALA, followed by GABA release may serve to coordinate the activity of GABAergic neurons in the nerve ring (e.g. SMDs) with that of ALA and AVE and eventually locomotory activity. While further studies are required to test the concept of “GABA recycling neurons” in *C. elegans*, we note an interesting precedent of GABA recycling in the vertebrate CNS. Midbrain dopaminergic neurons do not synthesize GABA, but take it up via the GABA transporters GAT1 and GAT4 and then release GABA to inhibit postsynaptic neurons (Tritsch et al., 2012; Tritsch et al., 2014).

We also discovered a group of unusual GABA-positive neurons, the AVA, AVB, AVJ and neurons (and possibly also male ray neurons). These cells express low but clearly detectable levels of GABA and require GAD for their GABA staining. However, these neurons fail to express the vesicular transporter UNC-47 or the GABA reuptake transporter SNF-11 (which, in other systems is sometimes used to release GABA, rather than reuptake GABA). Their GABA-positive nature does not depend on SNF-11 and these neurons therefore do not serve to clear GABA. Since they do not express known transporters to release GABA, it is not clear what purpose GABA serves in these neurons. Perhaps these neurons utilize novel means to employ GABA in synaptic signaling.

In conclusion, the extent to which GABA recycling or GABA clearance neurons exist in the vertebrate CNS remains unclear but we have used here the simplicity of the *C. elegans* nervous system to precisely define the set of GABA synthesizing and GABA reuptake neurons.

### GABA in non-neuronal cells

Apart from the easily explicable detection of GABA in muscle cells, the targets of the main class of GABAergic motor neurons, we detected GABA in two intriguing and unexpected non-neuronal cell types, the unusual hmc and the glia-like GLR cells. Both cells types may operate in GABA clearance. In vertebrates, some glial cell types are thought to employ GABA as a “gliotransmitter”, releasing GABA via a reversal of the plasma membrane GABA transporter GAT-1 to signal to neurons (Barakat and Bordey, 2002; Koch and Magnusson, 2009; Yoon and Lee, 2014). The GLR cells indeed express the *C. elegans* ortholog of the GAT-1 GABA transporter (SNF-11) and it will be intriguing to test whether the GLRs indeed also engages in active GABA signaling.

### Regulation of the GABA phenotype

We used the map of GABA-positive neurons as an entry point to study how neurons acquire their GABAergic phenotype. We built on previous work that implicated a few factors in controlling GABAergic features, extending the mutant analysis of these factors and identifying novel combinatorial codes of transcription factors that define GABAergic identity. We also identified factors that define the identity of GABA clearance and recycling neurons. Our works corroborates and significantly extends a number of previously developed themes and concepts:

#### (1) Combinatorial transcription factor codes

Transcription factors that specify the GABAergic phenotype act in neuron-type specific combinations (**Fig.11B**). Each GABAergic neuron type uses its own specific combination of regulators and, hence, there is no commonly employed inducer of GABAergic identity. This conclusion could already be derived from previous work (Hobert et al., 1999; Jin et al., 1994) and we confirm this conclusion here by defining the nature of several of the combinatorial transcription factor codes. Nevertheless, there is a notable reiterative use of two different regulators, *nhr-67* (RME, AVL, RIS) and *lim-6* (AVL, RIS, DVB) in specifying GABA identity in different cellular context. The neurons that are specified by *nhr-67* and *lim-6* are synaptically connected (White et al., 1986) and perhaps these factors may have a role in circuit assembly as well, as previously suggested for other “circuit-associated transcription factors” (Pereira et al., 2015).

#### (2) Preponderance of homeobox genes

The majority of regulators of neuronal identity (of GABA, but also Glu and ACh neurons) are encoded by homeobox genes. Those that are not (*nhr-67* and *elt-1*) cooperate with homeobox genes (**Fig.11B**). This is notable in light of the fact that only ∼10% of all transcription factors encoded by the *C. elegans* genome are of the homeodomain type. This observation argues that homeobox gene may have been recruited into neuronal specification early in evolution and that these homeobox-mediated blueprints then duplicated and diversified to generate more and more complex nervous systems.

#### (3) Phylogenetic conservation

Vertebrate GATA2/3 factors are postmitotic selectors of GABAergic identity in multiple distinct GABAergic neuron types (Achim et al., 2014; Joshi et al., 2009; Kala et al., 2009; Lahti et al., 2016; Yang et al., 2010). We found that its *C. elegans* ortholog *elt-1* also specifies GABAergic neuron identity, apparently in conjunction with the *unc-30/Pitx* gene. Remarkably, a population of GABAergic neurons in the CNS also co-expresses the mouse orthologs of *elt-1* and *unc-30* (Kala et al., 2009). All other factors we identified in *C. elegans* have vertebrate orthologs as well and according to the Allen Brain Atlas (Sunkin et al., 2013) are expressed in the adult CNS. It will need to be tested whether these orthologs may also have a role are expressed in and function in GABAergic neurons. Notably, however, the *C. elegans* ortholog of the Dlx genes, well characterized selectors of GABAergic identity in the anterior forebrain of the mouse (Achim et al., 2014), does not appear to be involved in GABAergic neuron differentiation.

#### (4) Coupling of GABAergic identity with other identity features

The decision of acquiring a GABA-positive phenotype is coupled to the acquisition of other terminal identity features. This is evidenced by the genetic removal of transcriptional regulators described here; such loss does not only result in the loss of GABAergic features, but also the loss of expression of other genes that define mature neuronal features, such as neuropeptides, ion channels, monoaminergic transmitter receptors and others. Transcription factors that control the expression of distinct terminal identity features have been termed “terminal selectors” (Hobert, 2008) and much of the data shown here supports the terminal selector concept. However, there are also exceptions: the *lim-6* LIM homeobox gene controls expression of *unc-25/GAD*, but not *unc-47/VGAT* (Hobert et al., 1999). A similar de-coupling of regulation of terminal identity features has been observed in the specification of the serotonergic neuron type NSM (Zhang et al., 2014) and in cholinergic command interneurons (Pereira et al., 2015). In the case of NSM, there appear to be partial redundancies with another homeobox gene (Zhang et al., 2014).

### A systems-wide regulatory map of neurotransmitter specification

All of the four conclusions derived here from our analysis of *C. elegans* GABA-positive neuron specification conform with similar conclusions derived from the analysis of the specification mechanisms for *C. elegans* cholinergic neurons (Pereira et al., 2015), glutamatergic neurons (Serrano-Saiz et al., 2013) and monoaminergic neurons (Doitsidou et al., 2013; Sze et al., 2002; Zhang et al., 2014; Zheng et al., 2005). The regulatory mechanisms for all these transmitter systems can be synthesized into a “regulatory map” of neurotransmitter specification, shown in **Fig.11B**. As shown in this figure, a view across different neurotransmitter systems illustrates that the activity of individual terminal selectors of neurotransmitter identities is not confined to specific neurotransmitter systems. For example, the *ceh-14* homeobox gene acts with different homeobox genes to specify GABA identity (ALA; this paper), glutamatergic identity (Serrano-Saiz et al., 2013) or cholinergic identity (Pereira et al., 2015). This reuse is remarkable if one considers that the four most re-employed transcription factors (*unc-3, unc-42, ceh-14, unc-86*) are involved in specifying the neurotransmitter identity of 46 of the 69 neuron classes for which a neurotransmitter regulatory is known (**Fig.11B**). We conclude that the system-wide view of neuronal specification, using distinct neurotransmitter systems, has revealed commonly operating principles of neuronal specification (Hobert, 2016a, b).

## MATERIALS AND METHODS

### Mutant strains

The *C. elegans* mutants strains used in this study were: *unc-47(e307), unc-25(e156), snf-11(ok156), unc-30(e191), nhr-67(ot795); otEx5999 [nhr-67 fosmid, pMG92(unc-47^prom^::mCHOPTI)], lim-6(nr2073), ceh-10(gm127), ceh-10(gm133)/hT2, tab-1(ot796), tab-1(gk753), tab-1(ok2198); tab-1(u271), ceh-14(ch3), ceh-17(np1), unc-4(e120), elt-1(ok1002) IV/nT1 [qIs51] (IV;V), unc-42(e419)*

### Transgenic reporter strains

The *unc-47, unc-46, gta-1* and *tab-1* fosmid reporter construct was generated using λ-Red-mediated recombineering in bacteria as previously described (Tursun et al., 2009). For the *unc-47, unc-46, unc-25* and *gta-1* fosmid reporter, an SL2 spliced, nuclear-localized mChOpti::H2B sequence was engineered right after the stop codon of the locus. For the *tab-1* fosmid reporter, an SL2 spliced, nuclear-localized YFP::H2B sequence was engineered right after the stop codon of the locus.

Wit the exception of *tab-1*, fosmid DNA were injected at 15 ng/µL into a *pha-1(e2123)* mutant strain with *pha-1(+)* as co-injection marker (Granato et al., 1994). The *tab-1* fosmid reporter DNA was injected at 15 ng/µL into *tab-1(gk753) otIs549* mutant strain with *rol-6(RF4)* as co-injection marker. Some of the resulting transgenes were chromosomally integrated. Resulting transgenes are: *otIs564* for *unc-47*, *otIs568* for *unc-46*, *otIs570* for *snf-11*, *otEx6746* for *gta-1*, *otEx6747* for *tab-1*.

The following reporter strains were generated for this study by injecting the PCR product from pPD95.75 plasmids containing the upstream region of the gene at 5 ng/µL into a *pha-1(e2123)* mutant strain with *pha-1(+)* as co-injection marker: *nlr-1^prom^::GFP(otIs527*, 150bp upstream the ATG*); unc-47^prom^::gfp (otIs509*, 300bp upstream the ATG with a deleted *unc-30* binding site to reduce expression in D-type motor neurons*); unc-25^prom^::mCHOPTI (otIs549*, 5.1kb upstream the 4^th^ exon*); unc-25^prom^::GFP(*otIs514, 7kb upstream the 6^th^ exon*); unc-46^prom^::GFP(otIs575, 234bp upstream the ATG)*.

For the GABA receptor reporters, the respective promoter region was cloned in front either of the GFP::unc-54-3’UTR or HcRed::unc-54-3’UTR. For *lgc-38*, 3.5kb upstream the 3^rd^ exon were used; for *lgc-37*, 5kb upstream the ATG (plasmid TU#920) was used; for *gab-1*, 3kb upstream the ATG were used (plasmid TU#921 as reported in (Topalidou and Chalfie, 2011)). The *lgc-38* reporter strain is from (Wenick and Hobert, 2004). *lgc-37* and *gab-1* reporter strains were made by injecting a PCR product at 5ng/µL into a *pha-1(e2123)* mutant strain with *pha-1(+)* as co-injection marker (resulting transgenes: *otEx6748* for *lgc-37* and *otEx6749* for *gab-1*).

The fosmid WRM0619bE05 (elt-1(+)) was injected in the mutant strain *elt-1(ok1002) IV/nT1 [qIs51] (IV;V)* with either *unc-47^prom^::GFP*(*otEx6751*) or *unc-47^prom^::mCHOPTI* (*otEx6750*) as co-injection marker. After lines were generated, worms carrying the array were singled. After three days, plates containing 100% worms with the array were isloated and used for subsequent analysis of *elt-1*

The following additional, and previously described neuronal markers were used in the study: *unc-47^prom^::GFP (oxIs12), unc-47^prom^::GFP (otIs348)*, *ser-4 ^prom^::GFP(adEx1616), ser-2b ^prom^::GFP(otEx536), avr-15 ^prom^::GFP(adEx1299), flp-22 ^prom^::GFP(ynIs50), kal-1 ^prom^::GFP(otIs33), flp-10 ^prom^::GFP(otIs92), wgIs354 [elt-1::TY1::EGFP::3xFLAG + unc-119(+)], rab-3::NLS::tagRFP(otIs355), cho-1^fosmid^::SL2::YFP::H2B (otIs354), eat-4^fosmid^::SL2::YFP::H2B (otIs388), cho-1^fosmid^::SL2::mCHOPTI::H2B (otIs544), eat-4^fosmid^::SL2::mCHOPTI::H2B (otIs518), wgIs395 [unc-30::TY1::EGFP::3xFLAG + unc-119(+)], lim-6 ^rescuing^ ^fragment^::GFP (otIs157), nhr-67^fosmid^::mChOPTI (otEx3362)*.

### Generation of the *nhr-67(ot795)* deletion allele

The *nhr-67* null allele *ot795* was generated by transposon excision (MosDEL) as previously described (Frokjaer-Jensen et al., 2010), using ttTi43980, a *Mos1* insertion in the first intron of *nhr-67* kindly provided by the NemaGENETAG Consortium. The resulting *nhr-67(ot795)* allele contains a ∼4.5 kb deletion, including the whole *nhr-67* coding region except for the 1st exon, as verified by PCR analysis and sequencing.

### GABA staining

A previously GABA staining protocol (McIntire et al., 1993b) was modified in the following manner. L4/young adult hermaphrodites or males were fixed for 15min (as opposed to 24h) at 4°C in PBS (137 mM NaCl, 2,7mM KCl, 10mM Na_2_HPO_4_, 2mM KH_2_PO_4_), 4% paraformaldehyde/ 2,5% glutaraldehyde fixative (as opposed to 4% paraformaldehyde/ 1% glutaraldehyde fixative). After being washed three to four times in PBS/0,5% Triton X-100, the worm were rocked gently for 18h at 37°C in a solution of 5% β-mercapto-ethanol, 1% Triton X-100 in 0,1M Tris-HCL(pH 7.5) (as opposed to 0,125M Tris-HCL(pH 6.9)). The worms were washed four times in 1% Triton X-100/0,1M Tris-HCL(pH7.5) and one time in 1mM CaCl_2_/1% Triton X-100/0,1M Tris-HCL(pH7.5). A worm pellet of 20-50 µL were shaken vigorously in 1mL of 1mM CaCl_2_/1% Triton X-100/0,1M Tris-HCL(pH7.5) and 1mg/mL of collagenase type IV (C5138, Sigma) for 30min. The worms were then washed three times in PBS/0,5% Triton X-100. An extra step was added in order to quench the autofluorescence due to the glutaraldehyde: the worms were incubated for one hour at 4°C in a freshly made solution of PBS and 1mg/mL of H_4_NaBo (Sigma, 71321).

Samples were blocked for 30 min at room temperature with 0.2% gelatine from fish (Sigma). Anti-GABA antibodies (abcam, ab17413) were used at a 1:250 dilution. For double labelling, anti-GFP (Thermo Fisher, A10262) or anti-RFP (MBL PM005) were used at a 1:1000 and 1:500 dilution respectivaly. Incubation were done overnight at 4°C. Secondary antibodies included Alexa-488-labelled-goat-anti-chicken (Invitrogen, A11039), Alexa-488-labelled-goat-anti-guinea pig (life, A11073), Alexa-555-labelled-goat-anti-guinea pig (life, A21435) or Alexa-594-labelled-donkey-anti-rabbit (Invitrogen, A21207).

### EMS screen and *tab-1* cloning

An Ethyl methanesulfonate (EMS) mutagenesis was performed on the reporter strain *otIs509* driving GFP expression in the 26 “classic GABA neurons”. 6762 haploid genomes were screened for abnormal expression of GFP. A mutant lacking *gfp* expression in RMEL/R (*ot796*) was isolated. After checking for the recessivity of the allele, *ot796* was crossed into the Hawaïan strain and 51 F2s missing RMEL/R were isolated and prepared for Whole genome sequencing as described in (Doitsidou et al., 2010). The results were then analyzed employing the CloudMap data analysis pipeline (Minevich et al., 2012). Complementation tests between *ot796* and three alleles of *tab-1 (u271, ok2198* and *gk753*) confirmed that *ot796* is an allele of *tab-1*.

### Microscopy

Worms were anesthetized using 100 mM of sodium azide (NaN_3_) and mounted on 5% agarose on glass slides. All images (except Fig.5 B,C,E, Fig.7, Fig.8B and Fig.9) were acquired using a Zeiss confocal microscope (LSM880). Several z-stack images (each ∼0.45 µm thick) were acquired with the ZEN software. Representative images are shown following orthogonal projection of 2–10 z-stacks. Images shown in Fig.5 B/C/E, Fig.7, Fig.8B and Fig.9 were taken using an automated fluorescence microscope (Zeiss, AXIOPlan 2). Acquisition of several z-stack images (each ∼0,5 µm thick) was performed with the Micro-Manager software (Version 3.1). Representative images are shown following max-projection of 2–10 z-stacks using the maximum intensity projection type. Image reconstruction was performed using ImageJ software (Schneider et al., 2012).

## ACKNOWLEDGEMENTS

We thank Q. Chen for generating transgenic strains, L. Carnell, B. Harfe, A. Fire and M. Chalfie for communicating unpublished results on *tab-1*, I. Topalidou and M. Chalfie for *gab-1* and *lgc-37* reporter plasmids, J. Ghergurovich for help with genetic screening and the original isolation of the *tab-1* allele, E. Serrano-Saiz and L. Pereira for advice on cell identifications and comments on the manuscript, C. Grove for the fly-over movie, J. Rand and J. Huang for discussions, S. Cook for analysis EM cross sections, the CGC for strains and M. Sarov for fosmid reporters. This work was funded by the National Institutes of Health [R01 NS039996] and the Howard Hughes Medical Institute. M.G. was supported by EMBO and HFSPO postdoctoral fellowships.

---

**Fig.5-figure supplement 1:**
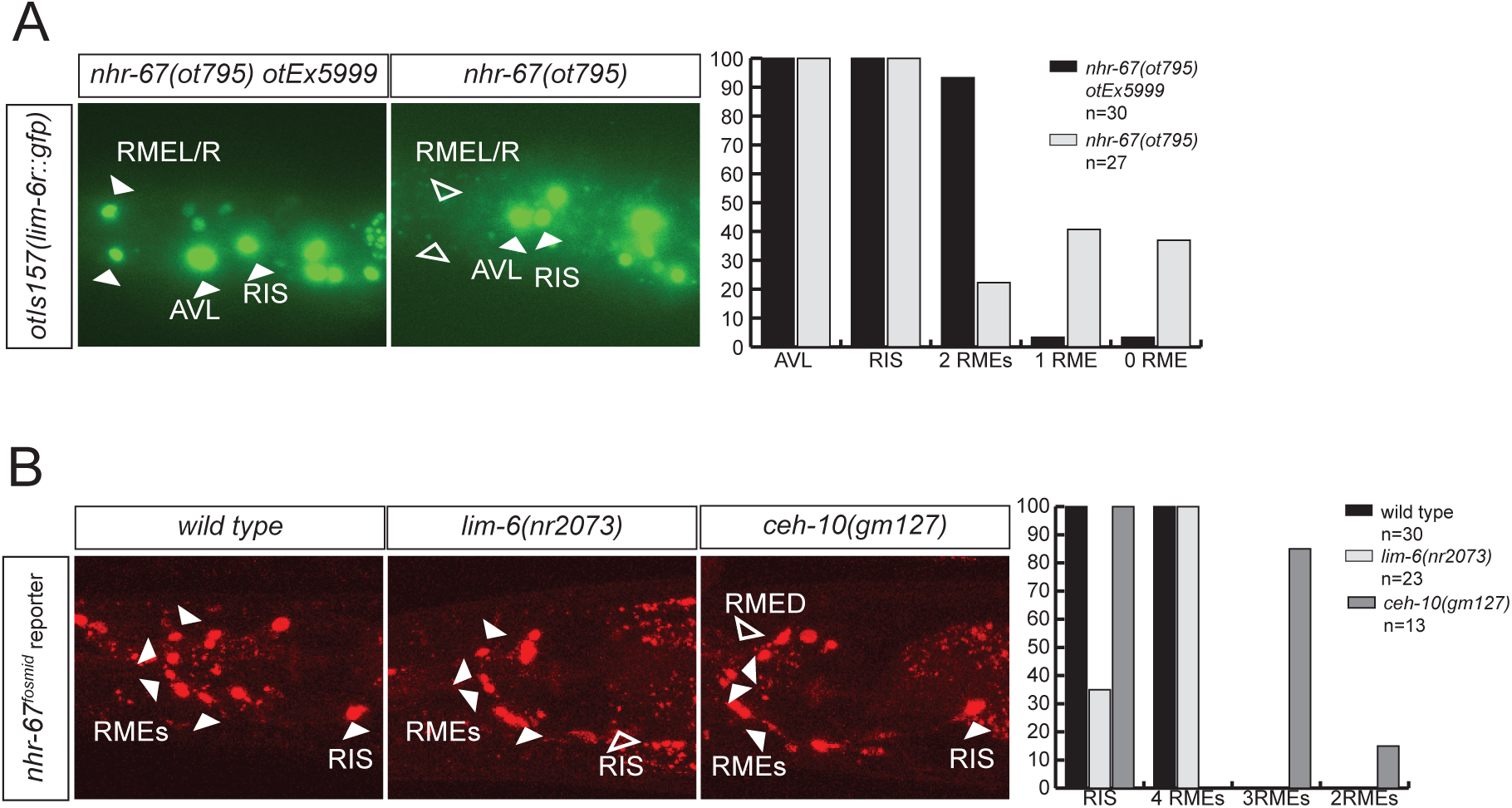
Regulatory relation of nhr-67 and other homeobox genes. **A:** *lim-6* expression is regulated *by nhr-67* in RMEL/R but not in RIS or AVL. **B:** *nhr-67* fosmid expression is controlled by *ceh-10* in RMED and by *lim-6* in RIS.

**Fig.6 – supplement 1:**
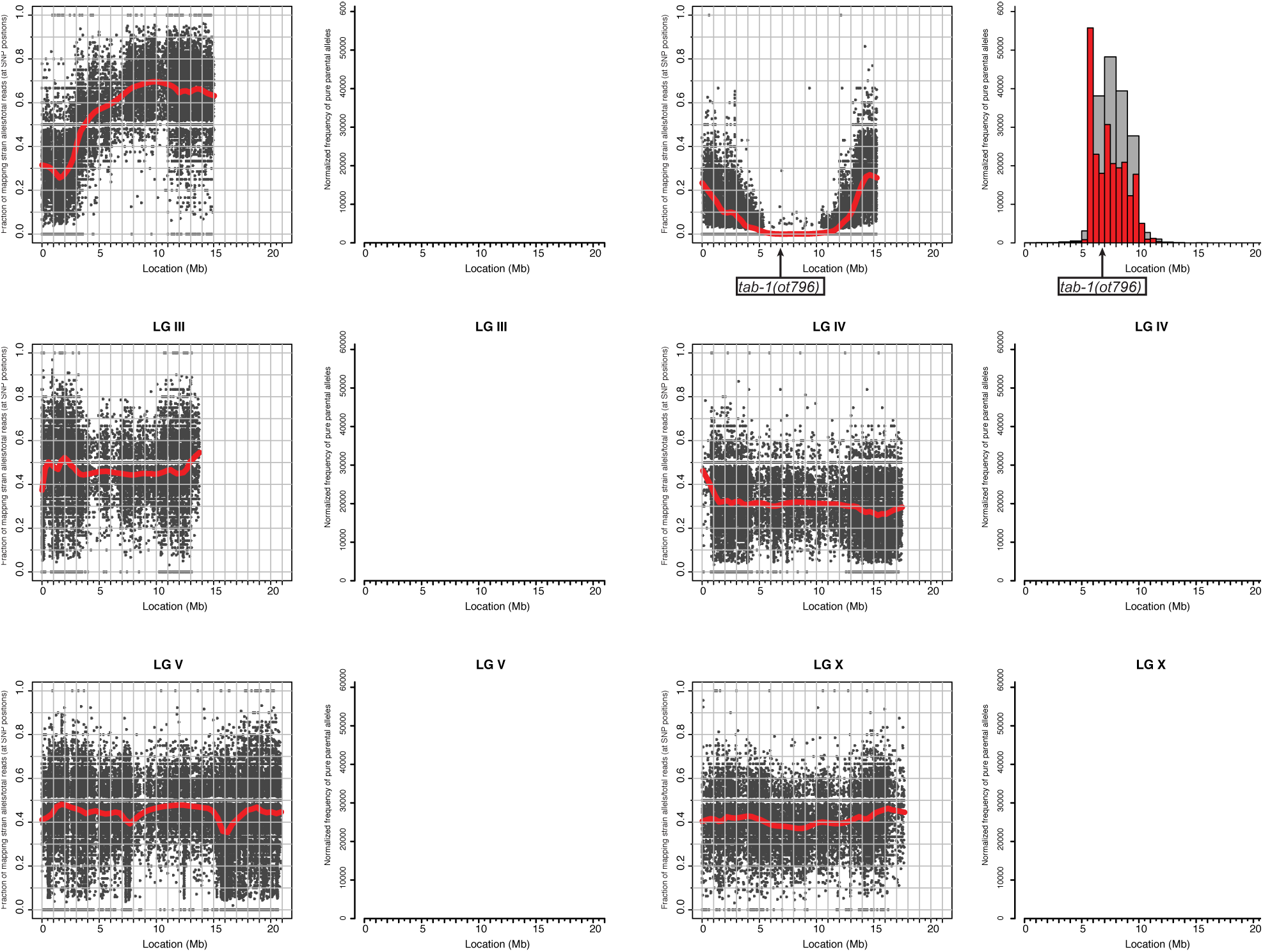
Whole genome sequencing data for *tab-1* mutant identification. A previously described mapping/sequencing approach was used (Doitsidou et al., 2010) and data was analyzed using CloudMap (Minevich et al., 2012).

**Movie 1: Flyover movies ACh (red), Glu (mint), GABA (blue), monoaminergic (green) neurons**. This movie was generated by Chris Grove.

